# A stimuli-responsive, pentapeptide, nanofiber hydrogel for tissue engineering

**DOI:** 10.1101/565317

**Authors:** James D. Tang, Cameron Mura, Kyle J. Lampe

## Abstract

Short peptides are uniquely versatile building blocks for self-assembly. Supramolecular peptide assemblies can be used to construct functional hydrogel biomaterials—an attractive approach for neural tissue engineering. Here, we report a new class of short, five-residue peptides that form hydrogels with nanofiber structures. Using rheology and spectroscopy, we describe how sequence variations, pH, and peptide concentration alter the mechanical properties of our pentapeptide hydrogels. We find that this class of seven unmodified peptides forms robust hydrogels from 0.2–20 kPa at low weight percent (less than 3 wt. %) in cell culture media, and undergoes shear-thinning and rapid self-healing. The peptides self-assemble into long fibrils with sequence-dependent fibrillar morphologies. These fibrils exhibit a unique twisted ribbon shape, as visualized by TEM and Cryo-EM imaging, with diameters in the low tens of nanometers and periodicities similar to amyloid fibrils. Experimental gelation behavior corroborates our molecular dynamics simulations, which demonstrate peptide assembly behavior, an increase in β-sheet content, and patterns of variation in solvent accessibility. Our Rapidly Assembling Pentapeptides for Injectable Delivery (RAPID) hydrogels are syringe-injectable and support cytocompatible encapsulation of oligodendrocyte progenitor cells (OPCs), as well as their proliferation and three-dimensional process extension. Furthermore, RAPID gels protect OPCs from mechanical membrane disruption and acute loss of viability when ejected from a syringe needle, highlighting the protective capability of the hydrogel as potential cell carriers for transplantation therapies. The tunable mechanical and structural properties of these supramolecular assemblies are shown to be permissive to cell expansion and remodeling, making this hydrogel system suitable as an injectable material for cell delivery and tissue engineering applications.

## Introduction

An attractive approach to tissue regeneration relies on water-swollen polymeric networks known as hydrogels, which can mimic features of the native extracellular matrix (ECM) such as proteolytic remodeling, cell-adhesion and mechanical properties, while guiding stem/progenitor cell fate decisions. Tissue engineering using biologically relevant hydrogel culture systems may improve regeneration, as hydrogels have a broad range of structural flexibility, biological activity, and similar mechanical properties as native tissue; this, in turn, can yield more physiologically-relevant cellular behavior^1–4^. However, there continues to be a need to develop suitable microenvironments that retain relevant biological and structural functions. Further progress requires materials that 1) are simple and inexpensive to synthesize, 2) gel in response to cytocompatible stimuli, such as small shifts in pH or temperature, and 3) support and regulate cell function, as a substitute for their normal physiological microenvironment^4^.

Transplanting stem cells may improve behavioral recovery following an injury or insult^5^, or during chronic or degenerative diseases^6^. However, transplant cell viability is often poor^6–8^, at least partly due to negative effects (cellular damage) that occur during injection, wherein cells undergo stresses such as non-physiological elongational flow and super-physiological shear forces^6^. Utilizing injectable and self-healing hydrogels as cell carriers could increase the surviving percentage of transplanted cells post-injection into damaged tissue for therapeutic repair^9^.

Oligopeptides are versatile building blocks that can be engineered to create self-assembled supramolecular structures which build upon non-covalent, reversible bonds^10–14^. For example, peptide amphiphiles spontaneously self-assemble into hydrogels in aqueous solution, driven by a sequence’s hydrophobic alkyl tail and a hydrophilic peptide domain^15^. Similarly, short amphiphilic peptides, such as LIVAGD^16^, form robust hydrogels via β-sheet assembly. Several other β-sheet forming peptides feature aromatic interactions (π-π stacking) that drive fibril formation and gelation, e.g., NFGAIL^16–18^ (a fragment of human islet amyloid polypeptide_22-27_), DFNKF^19–20^ (human calcitonin-derived peptide hCT_15-19_), and KLVFFAE^21^ (parts of amyloid β_16-22_). Recently, the 8-residue FDFSFDFS^22^ sequence and two pentapeptide analogs of an IDIDI^23^ sequence were shown to self-assemble into hydrogels. Likewise, a K_2_(QL)_6_K_2_ sequence also demonstrates self-assembly capabilities, where hydrogel formation is driven primarily by ionic screening of charges^13, 24^. Such self-assembling peptide materials can be designed to gel after injection, enabling uniform encapsulation of cells in 3D, ex vivo, and minimally invasive injection into target tissue such as brain and spinal cord^25–26^.

Several challenges limit the broader utility of these peptide systems in tissue engineering and regenerative medicine. First, protecting groups such as acetyl^27^, *t*-butyloxycarbonyl^28^, other large aromatic groups^29–30^, or the incorporation of D-stereoisomers^31^ are often required to induce gelation, thereby complicating synthetic processes and limiting scalability. Second, there are relatively few examples of short oligopeptides that exist beyond the derivation or analogs of the well-established diphenyl-alanine peptide sequence^32–33^. While these dipeptides form robust hydrogel systems, the hydrophobicity of the sequence limits their solubility and their range of potential applicability. Third, rather than having complete design freedom, many oligopeptides have been derived by sequence mapping onto relevant biological systems known to self-assemble into a variety of nanostructures. For example, designer peptide scaffolds have been based on an (EAK)_16_ sequence derived from the DNA-binding protein zuotin^34^.

We seek an approach to peptide-based hydrogel design that leverages the power of computation to guide peptide engineering efforts. Candidate peptides can be modeled in silico, using molecular dynamics (MD) simulations to interrogate, in atomic detail, the physicochemical properties of a given sequence^35–39^. Few examples exist of using computational approaches to design functional peptide scaffolds for tissue regeneration applications. The complex interactions between polypeptides and their environments, which mediate their self-assembly into useful biomaterials, demand new tools for characterizing the structure and function of peptide ECMs. An integrated approach that leverages computational modeling to understand experimental peptide behavior can more rapidly (and affordably) examine assembly. Moreover, such an approach could even survey the suitability of an array of potential sequence constructs (i.e., a library) toward creation of a general-purpose hydrogel scaffold for tissue engineering applications.

Here, we use an integrated computational and experimental approach in the design, synthesis, and characterization of a pH-triggered, self-assembling pentapeptide suitable as a three-dimensional scaffold for cell culturing. We term these materials *Rapidly Assembling Pentapeptides for Injectable Delivery* (RAPID) hydrogels. To decipher the self-assembly mechanism, we analyzed the peptide sequence KYFIL, with the goal of identifying specific residues that play a key role in intermolecular association and self-assembly. We screened seven different sequences for their ability to form hydrogels, and analyzed their supramolecular assembly behavior using a host of methods: attenuated total reflectance–Fourier-transform infrared (ATR-FTIR) spectroscopy, cryogenic electron microscopy (cryo-EM) and transmission electron microscopy (TEM), rheometry, and molecular dynamics (MD) simulations. RAPID hydrogels were also evaluated as three-dimensional scaffolds for cell encapsulation, via cell culture–based biological assays, thus allowing us to determine their cytocompatibility under physiological conditions.

While demonstrating a wide range of stiffnesses suitable to emulate a variety of human tissues, we focused here on tailoring RAPID hydrogels to mimic the biomechanical properties of brain tissue (Young’s moduli ranging 0.1–3.5 kPa^1, 40^). Encapsulation of oligodendrocyte progenitor cells (OPCs) enabled us to demonstrate the 3D cell culture matrix potential of RAPID hydrogels. We find that RAPID hydrogels can also mitigate the damaging effects of extensional flow experienced by cells during syringe injections.

## Results and Discussion

We have designed pentapeptides, based on a KYFIL-NH_2_ sequence (Figure 1) hereafter referred to simply as ‘KYFIL,’ that can self-assemble into β-sheet–forming nanofibers. The sequence KYFIL was chosen based on previously published results on aromatic-rich tripeptides that could gel under certain experimental conditions^27, 30, 35, 41–42^ (such as a change in pH or ionic strength). In particular, we chose Lys as the head-group to improve solubility in aqueous solution^24^, while the overall sequence design was guided by the goal of increasing the hydrophobicity of amino acid residues so as to increase the amphiphilicity of the peptides. In initial screens, we assayed several peptide designs by performing an alanine scan. Displaying the results as a sequence logo^43–44^ revealed that the central phenylalanine (F), as well as preservation of the amphiphilicity of the sequence, are two key elements that facilitate hydrogel formation (Figures 1 and 2). Interestingly, the carboxyl-terminated variant KYFIL-CO_2_H (i.e., with ‘natural’ peptide end-chemistry) did not readily form hydrogels at pH 7.4. Because the carboxylic acid moiety is deprotonated at neutral pH, this finding suggests that an uncharged C-terminus is required for gelation of the peptide. By examining the peptide’s secondary structural conformation via ATR-FTIR spectroscopy and MD simulations, we detected that structural transitions occur when a pentapeptide self-assembles under gelling conditions.

**Figure 1.**
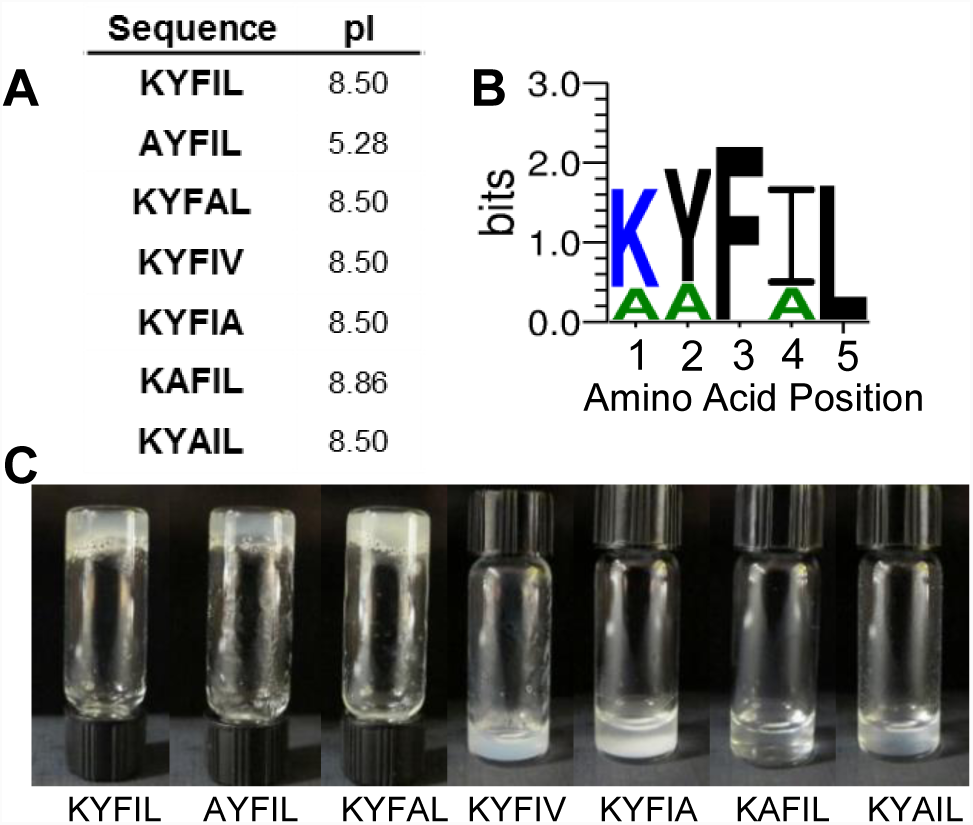
Sequences investigated in this study. A) Each sequence examined is listed, along with its theoretical isoelectric point (pI). All peptides are C-terminally amidated. B) A sequence logo highlights the order and predominance of amino acids within pentapeptide analogs that gel under any pH condition. The sequence profiling suggests that Phe (F) and Leu (L) must be conserved for gelation. C) When peptides are dissolved in PBS at pH 7.4 and 1.5 wt. %, the KYFIL, AYFIL and KYFAL pentapeptides form hydrogels, whereas other sequences do not gel under these conditions; note that KAFIL can gel at pH > 10.

**Figure 2.**
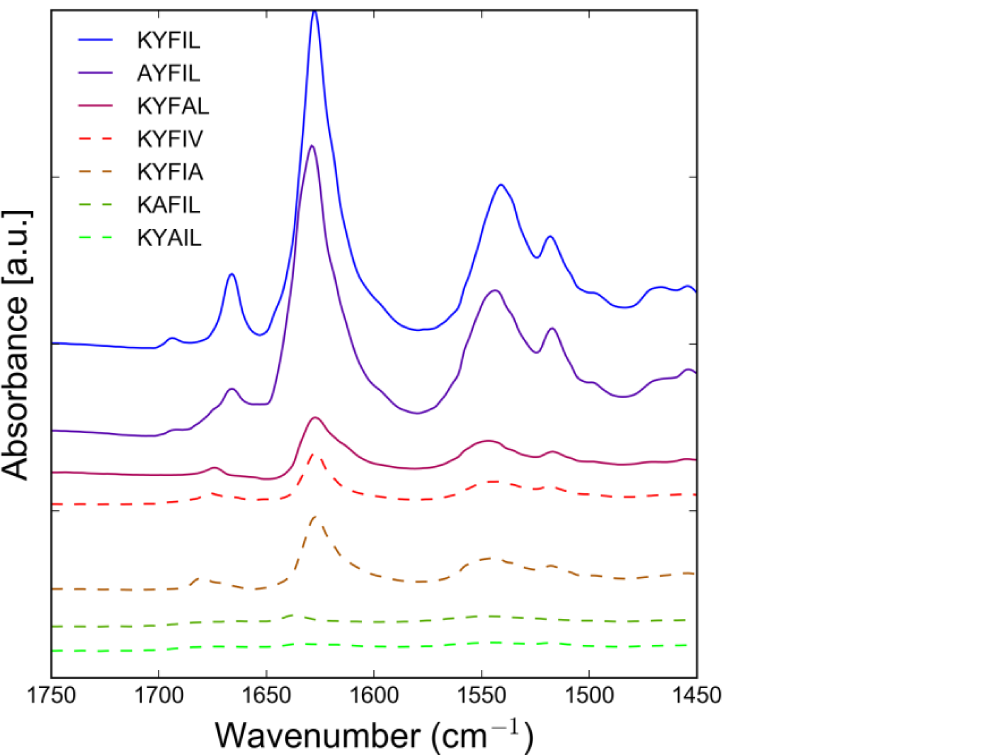
Peptides exhibit characteristic secondary structures via ATR-FTIR. Peptides dissolved at 3 wt.% in PBS and pH 7.4 were examined. All gelling peptides (solid lines) exhibit an Amide I absorbance at 1629 cm^−1^, indicative of β-sheet hydrogen bonding. A peak near 1679 cm^−1^ to 1683 cm^−1^ suggests anti-parallel β-sheet conformation. Non-gelling peptides (dashed lines) exhibit much weaker, less intense peaks at the same wavenumbers. All spectra are baseline corrected, normalized, and vertically offset for clarity.

### Secondary Structure Analysis via FTIR Spectroscopy

The secondary structural content of pentapeptide specimens was probed by ATR-FTIR spectroscopy. Samples of the pentapeptides of interest were generated via solid-phase peptide synthesis methods. An alanine (Ala) scan^45^ of KYFIL was used to assess each AA’s contribution to gelation, wherein individual amino acids (AAs) were sequentially exchanged with Ala or Val. Ala and Val were the substituted residues of choice, as they either eliminate a side-chain beyond the Cβ atom (Ala) or otherwise would be expected to minimally alter the main-chain conformation (Val). As uncharged and relatively compact residues, Ala and Val would not be expected to introduce (confounding) electrostatic or steric effects^46^. All pentapeptides were dissolved in phosphate-buffered saline (PBS) at a concentration of 3% (weight/volume) and adjusted to a pH of 7.4. Gelling peptides (solid lines, Figure 2) display a strong Amide I absorbance at ≈1629 cm^−1^, arising from vibrational modes of the amide group; these vibrations, in the region of 1700→1600 cm^−1^, correspond to stretching of the C=O and C–N bonds, as well as bending of the N–H^47–48^. This region of the IR spectrum is particularly sensitive to variations in secondary structural conformation and, in the case of our pentapeptide samples, is indicative of β-sheet hydrogen bonding^49–50^ (Figure 2, Figure S2 and S3). A secondary peak near 1679 cm^−1^ to 1683 cm^−1^, in some of the specimens, indicates that the β-sheet is anti-parallel^51^. We can infer this because the Amide I region of parallel β-sheets harbors a single predominant signature (near 1630 cm^−1^), while antiparallel sheets generally feature a second (minor) peak near ≈ 1680-1690 cm^−1^. Peptide variants which do not form gels at the same concentration and pH (dashed lines, Figure 2) exhibit less intense peaks, suggesting a lack of significantly structured hydrogen bonding networks in those solutions. By correlating our IR observations with variations in AA sequence, it appears that amphiphilicity and a capacity for π-system interactions (e.g., π…π stacking and π…cation interactions with the benzyl side-chain of the central Phe) play a key role in self-assembly and hydrogel formation (Figure 2).

### Probing the Conformational Space and Interaction Events via MD Simulations

MD simulations were used to examine the atomically-detailed molecular interactions underlying peptide self-assembly processes. MD simulations offer a powerful approach to examine the structural properties and conformational dynamics of engineered peptides and can yield experimentally-inaccessible insight about the dynamical basis of self-assembly^52–53^. Simulations can help guide adjustments to the peptide sequence in order to optimize the system’s properties toward a target goal. Using MD simulations, one can study a peptide system’s aggregation propensity by simulating multiple peptides together in a single system. While other work has focused on the diffusional association of protein molecules within a solvated system^39, 54^ or detailed the biomolecular recognition events (i.e. conformational rearrangement and binding/unbinding events^55^), few examples exist of using computational approaches to design functional peptide scaffolds for tissue regeneration applications^56^. In this study, we used MD simulations to study the emergence of structural features in a peptide system and provide an atomistic view of the self-assembly process of nanostructures.

To elucidate the molecular-scale events associated with self-assembly, after having experimentally established pentapeptide sequences that can assemble in aqueous media, we conducted MD simulations of select peptide candidates (Figure 3). These extended (200-ns), all-atom simulations were performed in explicit solvent using the CHARMM36 force-field. Such force-fields represent the physicochemical properties of each amino acid—including partial charges, atomic interaction (Lennard-Jones) potentials, and other parameters—via a classical, molecular mechanics–based approach, as described in various primers^52^. In practice, CHARMM36^57^ is a state-of-the-art force-field that can be applied to many types of biomolecular systems, as illustrated for instance by the analysis of disordered regions of the protein desmoplakin^58^. Our simulations show that the assembly propensity of RAPID peptides correlates with the diffusional association of individual peptides. The peptides primarily adopt irregular conformations, with some transiently-stable β-turns ‘flickering’ into existence (Movie S1–S5). Within ≈50 nsec, individual KYFIL peptides assemble into six discrete groups of peptides, as can be seen by visual inspection of trajectories, with some β-sheet secondary structure (Figure S1 and Figure 3A). Extending the simulation further yields peptides that have assembled into two large clusters by ≈200 nsec (Figure 3A).

**Figure 3.**
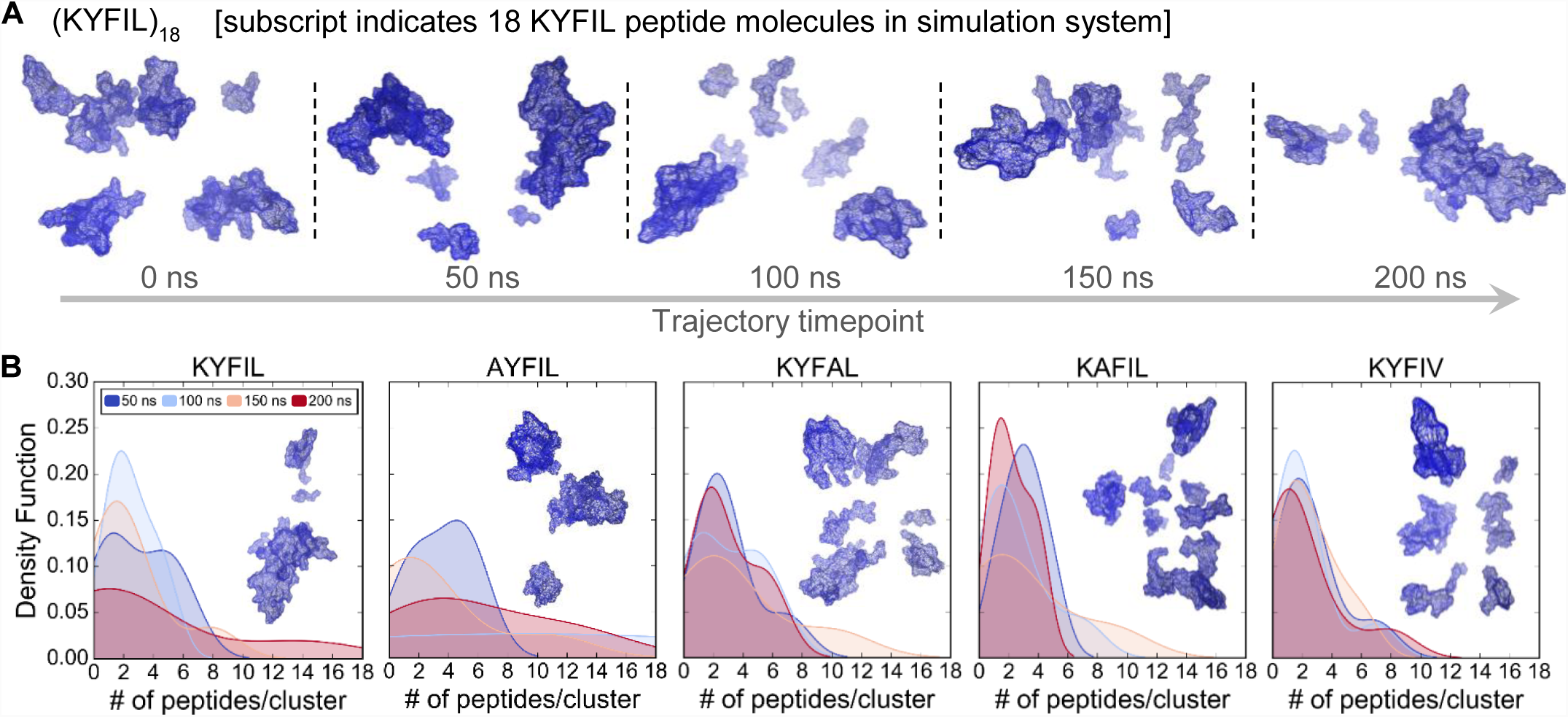
KYFIL peptide molecules simulated in explicit solvent assemble into multimeric structures. A) Representative structures of the simulated pentapeptide sequence, KYFIL. Spatiotemporal evolution of peptide assembly is demonstrated from the simulation trajectory of the peptides at 50 ns intervals as the molecules assemble into large clusters on the timescale of 200 ns. B) Density functions representing the clustering propensity of different pentapeptide systems over time. At the end of the 200 ns simulation, KYFIL has the least number of distinct clusters and largest number of peptides per cluster, versus other peptide sequences. Insets are representative snapshots of the peptides near 200 ns.

We used a discrete number-density function (a measure of the local concentration) to quantify the aggregation propensity of the four different peptides along their respective trajectories. Those sequences which were found to gel in experiments— KYFIL, AYFIL, KYFAL, and KAFIL at pH 10—exhibited a higher propensity to aggregate; those peptides which formed gels also tended to exhibit greater variation in the number of peptides per cluster at 200 ns, consistent with a higher propensity to assemble, even non-specifically into heterogeneous aggregates (Figure 3B). In addition, the time-evolution of the radius of gyration (*R*_g_) of the peptide systems (*R*_g_ computed system wide across all peptides, not per-peptide) reveals a gross structural rearrangement—from mostly diffuse peptides to closely associated molecular interactions, as indicated by the net decrease in *R*_g_ for pentapeptide sequences (Figure S4) relative to the initial trajectory, except for KAFIL, KYFAL, and KYFIV. These data are consistent with the FTIR spectra (Figure 2), as KAFIL, KYFAL and KYFIV have lower β-sheet peak intensities, suggesting lower assembly propensity. For KYFIL, a detectable, and presumably hydrophobically-driven, ‘collapse’ of the system appears to be more kinetically allowed, versus other sequences; i.e. transitions between secondary structures occur frequently, implying relatively low activation barriers^59^. Visual inspection of trajectories shows a sharp structural reorganization early on (< 100 ns) in most of the simulations.

We also examined the structural transitions from the initial peptide system (post-equilibration) to the final conformational ensemble. For all simulated peptide sequences, there was a notable dearth of α-helicity (Figure 4A), consistent with the experimental FTIR data (Figure 2, Figure S3). All pentapeptides preferentially sampled β-type structures (Figure 4a), and the gelling peptide sequences (KYFIL, AYFIL, KYFAL, KAFIL) exhibited a nominally greater fraction of β-strand character over the course of the 200-ns trajectory, versus a non-gelling sequence, KYFIV (Figure 4B). The domain-swapping mode of β-rich association can be induced by intermolecular β…β-strand/bridge contacts, via directional hydrogen bonding between the backbones of aromatic residues and β-branched amino acids (e.g. isoleucine)^60–61^. Consequently, the structural rearrangement of peptides can reduce conformational strain, as the formation of such β-strand structures are enthalpically favorable, driving the folding of β-sheets^62–63^. The torsion angles for each type of amino acid, barring the N- and C-termini (Figure S5), indicate significant structural heterogeneity for each peptide system. Our results suggest that, in general, the middle Phe in each pentapeptide often adopts a type-II β-turn conformation (ϕ = −60°, φ = 120°) or an antiparallel β-sheet structure (ϕ = −140°, φ = 135°); this is consistent with our aforementioned FTIR results. For the KYFAL and KAFIL sequences, the Ala preferentially samples a polyproline type-II helix (ϕ = −75°, φ = 145°), with a decreased β-sheet propensity. This result is unsurprising, as the peptide backbone near an Ala (versus Tyr) residue encounters less steric hindrance, given the absence of the phenol side-chain^64^. The most densely populated regions of conformational space for Ile, in all pentapeptide sequences (Figure S5), highlights this amino acid’s propensity to adopt β-sheet conformations. In this context, Phe…Ile intermolecular interactions (steric occlusion, as well as London dispersion forces and other van der Waals forces) are particularly relevant, as they would facilitate the hydrophobic aggregation of these peptide regions and indirectly enable the formation of hydrogen-bond networks between the local backbones^65^; this model is also consistent with both MD simulations and FTIR spectroscopic data (Figure 2).

**Figure 4.**
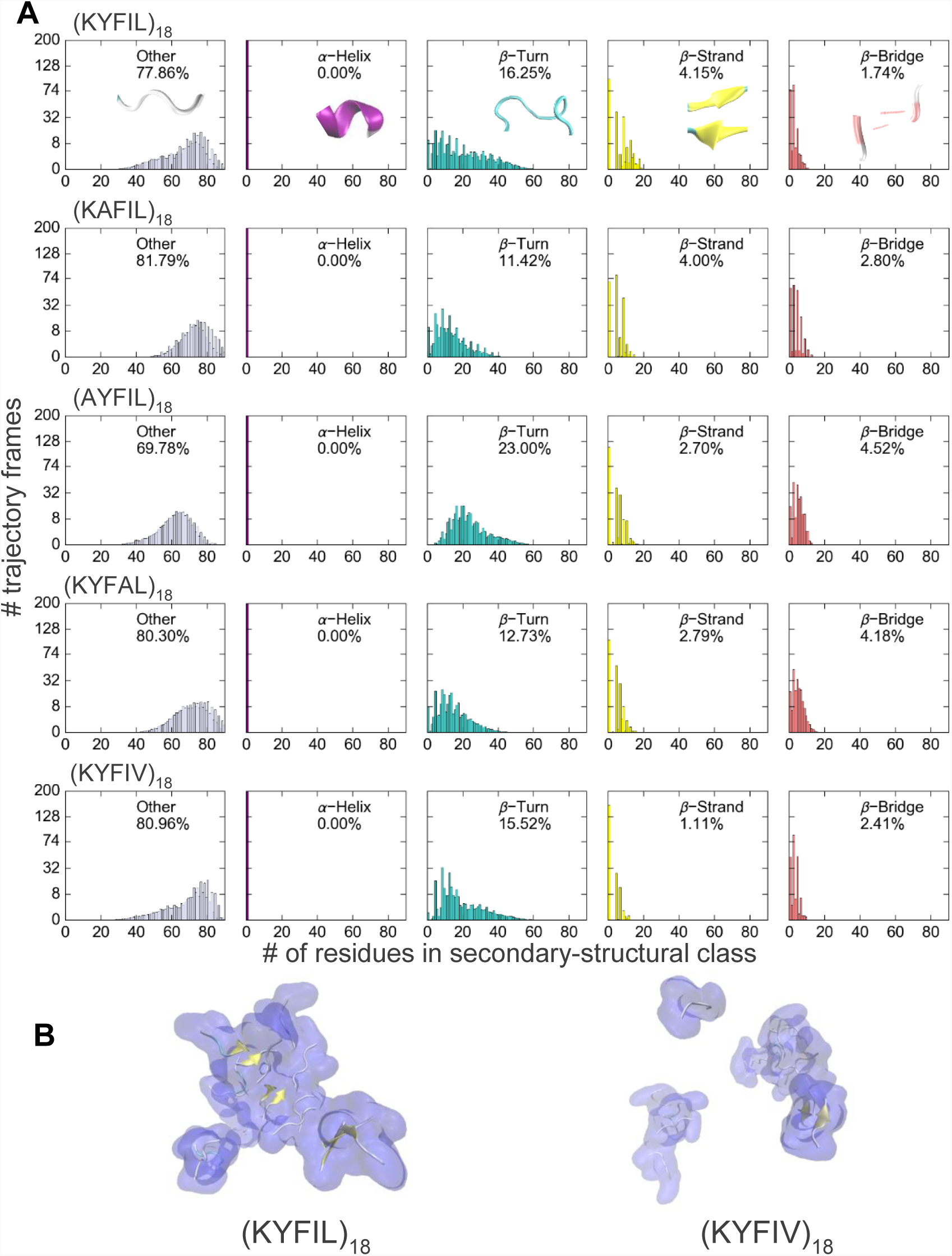
Secondary structural content of the simulated KYFIL, KAFIL, AYFIL, KYFAL, and KYFIV systems. Numbers within a plot represent the population of the secondary structure observed in the simulation over all possible secondary structure conformations. Secondary structure cartoon representations in the thumbnails displayed in the first row match the colors in the histogram. A) Histograms that depict the predominant conformations exhibited by the polypeptide are *β*-turns and ‘other’ structures. For all sequences, there is an absence of α-helical structures, consistent with our experimental results. In addition, *β*-turn structures are prevalent with *β*-strand and bridge structures. A significant shift from strand to bridge occurs in the character of the *β* structure in the non-gelling sequence, KYFIV. B) Representative snapshots taken at 180 ns and 177 ns for KYFIL and KYFIV, respectively, illustrating sequence-dependent conformational states of the pentapeptides. The peptides can be seen to be a mixture of helices and coils; the secondary structures are labelled in this view with α-helices colored purple, 3_10_ helices blue, *β*-strands yellow, the *β*-turn motif cyan and irregular coil regions white. These trajectory frames illustrate the formation of *β*-sheet regions within the two peptide systems, with more pronounced populations of *β*-sheet conformations present in KYFIL versus KYFIV.

In addition to internal (intra-peptide) and external (inter-peptide) interactions, the conformational dynamics of a peptide system are governed by peptide…solvent interactions. By quantifying peptide…water contacts, we can discern hydrophobic side-chain contributions to the energetics of peptide assembly, and also study a peptide’s solvation dynamics. Thus, we evaluated the solvent-accessible surface area (SASA) of individual residues in each pentapeptide, averaged over entire 200-ns trajectories. In computing the relative SASA of a peptide system (via Rost & Sander’s method^66^), we consider the ordinary accessibility of a residue in a structure normalized by the maximal value possible for that residue type (i.e., that amino acid sidechain). Unsurprisingly, for each pentapeptide sequence the N-terminal Lys was the most solvent-exposed (Figure 5) and the central tripeptide (…Tyr/Ala–Phe–Ile/Ala…) was the most consistently buried throughout the simulation. The significant changes in relative accessibility of the C-terminal residue (position 5) indicate the system’s structural rearrangement in the context of side-chain functional groups^67^. Our findings are consistent with patterns in Kyte-Doolittle hydropathicities^68^ (GRAVY values in Figure 5) as well as prior experimental results regarding the hydration structure of ABA triblock copolymeric systems^13, 24, 69–70^, wherein the termini were found to be exposed to aqueous solvent molecules and dehydration of non-polar side-chains biases the middle block of the amphiphilic pentapeptide to preferentially adopt compact 3D structures (β-strands, turns, etc.) that occlude solvent.

**Figure 5.**
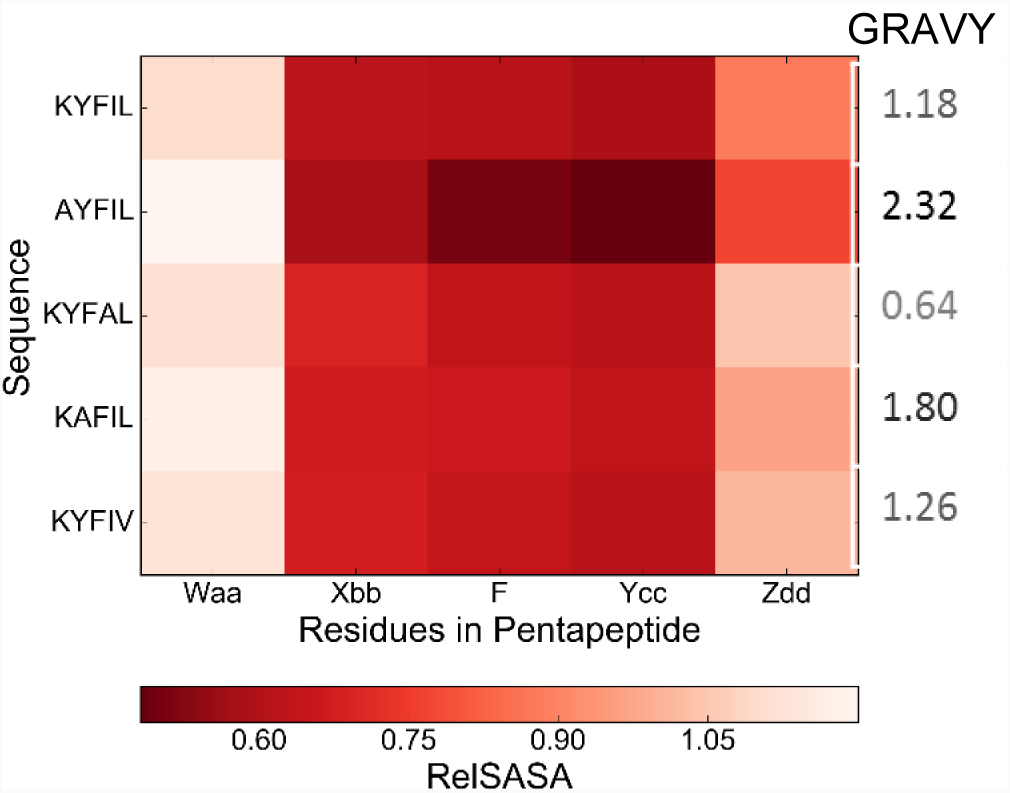
Sequence-dependent changes in relative solvent accessible surface areas (RelSASA) for individual residues in each pentapeptide simulation. The RelSASA quantifies the accessible surface area of each residue in the folded pentapeptide. A white color indicates that a residue is more solvent-exposed than average, while the intensity of a red scales with residue burial. Computed grand average hydropathicity (GRAVY) values, which are essentially Kyte-Doolittle (KD) hydrophobicity indices averaged over the amino acid sequence for each peptide, are given on the right; on the KD scale, the hydrophobic amino acids have positive values (the most hydrophobic is Ile, with a value of +4.5), while hydrophilic residues have negative values (the least hydrophobic is Arg, at −4.5, followed by Lys at −3.9). At least qualitatively, the MD-based results and general hydropathicity patterns are consistent: the most hydrophobic peptide, AYFIL (most positive GRAVY score), features the least solvent exposure over the course of its MD trajectory, while the most hydrophilic peptide, KYFAL (least positive GRAVY score), exhibits the largest RelSASA values.

### Hierarchical Self-Assembly: Evaluating Hydrogel Rheological Properties

The mechanical properties of 1.5 and 3 wt. % hydrogels were found to depend on the concentration, pH, and peptide sequence. Hydrogels formed *in situ*, in an epitube, within several seconds (Movie S6) and were then pipetted onto the rheometer platform for rheological measurements. RAPID hydrogel stiffnesses span two orders of magnitude, from approximately 50 – 17,000 Pa in shear storage moduli (G′) (Figure 6a). (For comparison, this would be similar to 520 – 44,200 Young’s modulus, although this requires a potentially fraught assumption of a Poisson’s ratio of 0.5.) KYFIL at 1.5 and 3 wt. % forms hydrogels of 8,000 and 17,000 Pa, respectively (Figure 6 and S6).

**Figure 6.**
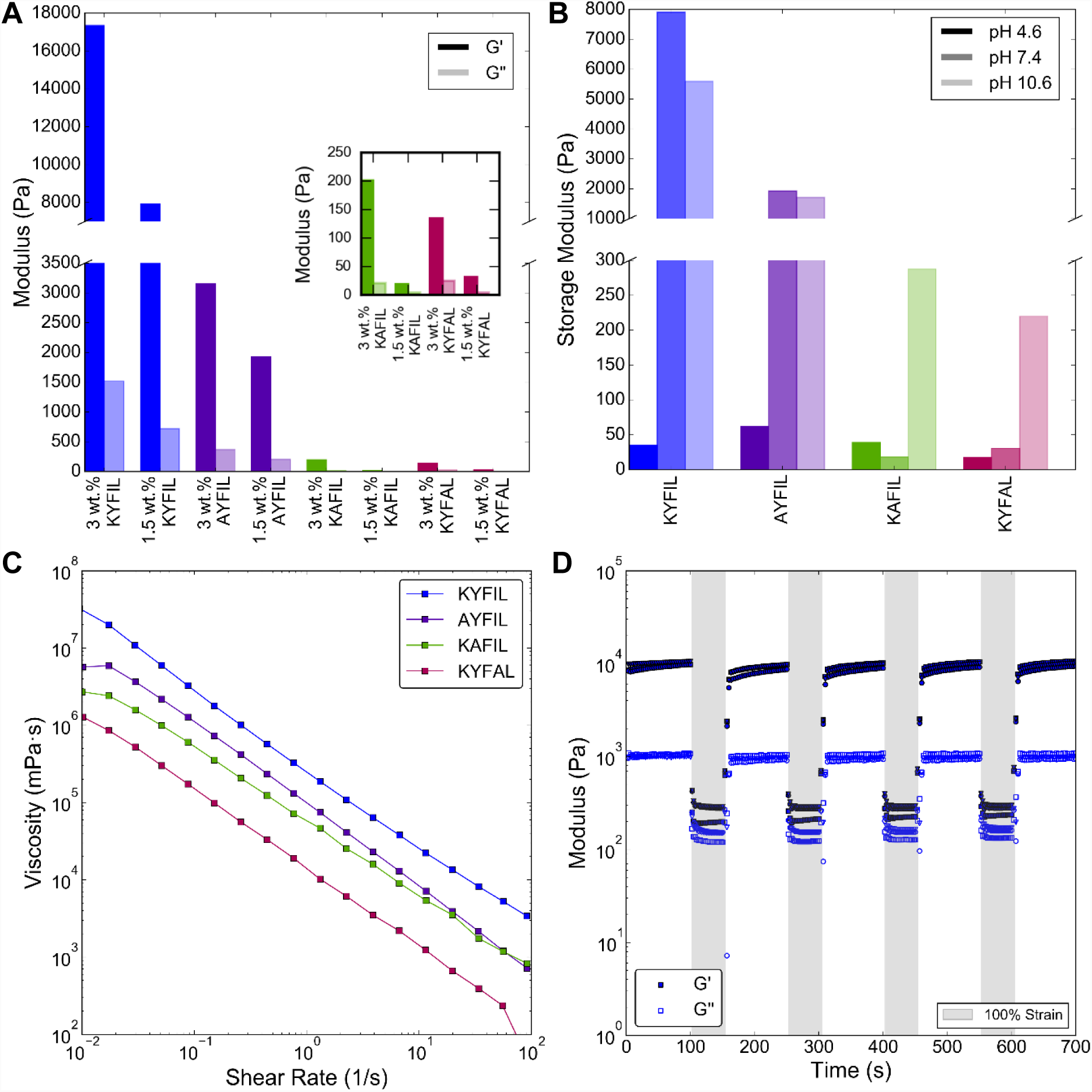
Rheological properties of self-assembling pentapeptides at different concentrations and pH conditions. a) Storage and loss moduli as determined from the linear viscoelastic region (LVE) taken from strain sweeps at a constant frequency of 1 hz of 1.5 wt. % and 3 wt. % hydrogels at pH 7.4. Hydrogels were formed *in situ* in an epitube and then pipetted onto the rheometer platform. Hydrogel stiffness can be tuned by concentration and peptide sequence variation. The inset is a magnification of the G’ and G” for KAFIL and KYFAL hydrogels. b) Storage moduli taken from the LVE from strain sweeps at a constant frequency of 1 hz of 1.5 wt. % hydrogels at different pH conditions of 4.6, 7.4, and 10.6. The mechanical properties of the hydrogel are dependent on pH, where all peptide sequences are very weak gels (G’ < 80 Pa) in acidic conditions, and form robust hydrogels at pH 7.4 and 10.6. c) Hydrogel forming sequences were evaluated under shear flow to determine their shear-thinning properties. The apparent viscosity of each sample decreased with increasing shear rate demonstrating that these hydrogels are capable of shear-thinning. d) 1.5 wt. % KYFIL hydrogels (n = 3) were subjected to five step strain sweeps of 100 % strain (50 s), followed by a 100 s recovery period (0.1 % strain). The hydrogel recovers 70-80% of its initial G’ within several seconds. Even after multiple high strain cycles, the hydrogel is able to repeatedly retain its mechanical strength.

Peptide hydrogels can provide structural flexibility and mechanical properties that emulate native biological tissues^4, 71–72^. Bulk matrix stiffness and topography are well known biomechanical cues that can direct stem cell proliferation as well as differentiation^73–74^. In most tissues, such as the heart, muscle and bone, the extracellular matrix contributes to the biophysical microenvironment, e.g. a Young’s modulus of 6.8 kPa for heart tissue and up to 103 kPa for bone^75–76^. However, tissues within the central nervous system (CNS), such as the brain and spinal cord, are some of the most compliant tissues in the body^77^, with moduli of ∼0.7 kPa to 3.5 kPa^78–80^. Such an extensive range of stiffness requires hydrogel biomaterials to have highly tunable biomechanical properties that can be catered to a wide range of applications, for numerous different tissue types throughout the body.

Our class of peptide sequences is unique in that the peptide lengths are quite short (5 amino-acid residues), and have a broad range of mechanical properties (∼50 – 17,000 Pa) that can be fine-tuned via small changes in concentration or pH (Figure 6). The broad range and large magnitude of storage moduli we can attain is in contrast to other short, self-assembling oligopeptides. For example, K_2_(QL)_6_K_2_, RADA16-I, (FKFE) _2_^81^, MAX1/8^82^, and KLVFF^83^ sequences yield gels with much lower storage moduli and narrower ranges of mechanical properties (storage moduli of 50 - 1000 Pa).^13, 84–85^

The lower storage modulus of KYFAL can be reconciled with its weaker signature peak intensities for β-sheets in the FTIR spectra (Figure 2), implying less content of well-ordered β-sheet for KYFAL (Figure 4). Additionally, Lys did not seem to affect gelation, so long as the amphiphilicity of the sequence was maintained. Rather, the substitution of Lys → Ala affected the solubility of the peptide (Figure S8). Similarly, at 3 wt. %, KAFIL had a G′ of 200 Pa compared to KYFAL at 133 Pa. The additional bulky methyl group in the Ile–Leu C-terminus of KAFIL, relative to Ala–Leu (in KYFAL) or Ile–Ala and Ile– Val (in KYFIA and KYFIV, respectively) confers greater hydrophobicity, resulting in a self-assembly process driven mainly by increased hydrophobic interactions.

The apparent pKa shift following the substitution of Tyr to Ala increases the electrostatic repulsion between peptides^30^, reducing the aggregation propensity. Increasing the pH, which alters the average degree of ionization, better neutralizes KAFIL and favors self-assembly of the peptide, by reducing the mean net charge of the Lys head group^86^. Our rheological characterization of hydrogel-forming peptides indicates that Ile facilitates self-assembly^87^: we detect a higher population of β-sheet conformations for Phe–Ile–Leu versus Phe–Ala–Leu sequences.

In investigating the pH responsiveness of hydrogel-forming sequences, we found that all peptides exhibited lower storage moduli upon a decrease in pH. The three pHs were chosen (4.6, 7.4 and 10.6) to include the physiological pH of 7.4—particularly relevant for viable cell encapsulation—as well as acidic and basic pHs that bracket the pI of each sequence (Figure 1A). The G′ increases by several orders of magnitude as the pH of the solution increases toward neutrality (Figure 6B). Non-gelling sequences (KYFIV, KYFIA, KYAIL) also exhibit pH-responsive behavior: at low pH, the peptides were soluble, but precipitated as an off-white powder as the pH was raised (but never gelled). The storage (G′) and loss (G″) moduli of 1.5 and 3 wt. % hydrogels increased with increasing concentrations of the hydrogel and increasing pH conditions (Figure 6A, B, S7 and S8).

The apparent viscosity of all gelling sequences decreased linearly with increasing shear rate, demonstrating the shear-thinning capacity of these hydrogels (Figure 6C, Figure S10). Multiple high-strain (100%) sweep cycles, with 30 s recovery periods, demonstrated KYFIL’s ability to self-heal following mechanical deformation, without any evidence of hysteresis (Figure 6d, Figure S11). Following a 100% strain, hydrogels repeatedly recovered gel behavior within 14 seconds (G’ > G”). Within 1 minute, the gel recovered 82% of its initial G’, and required 3.4 minutes to recover 90% and 7 minutes to recover 96% (Figure S11). Even after multiple high-strain cycles, the hydrogel rapidly and repeatedly recovers its mechanical strength—rendering these materials particularly ideal for biomedical applications that require injection. This enables uniform encapsulation of cells in 3D, *ex vivo*, and then injection via a minimally invasive technique. Similarly, we found that the hydrogels could re-gel, macroscopically, following a syringe ejection (Movie S7), suggesting that materials based upon these peptides could be well-suited to additive manufacturing applications like extrusion-based 3D printing.

The propensity of our RAPID peptides to adopt β-rich structures, alongside their capacity to form hydrogels (and the presence of fibrillar networks in such gels [see below]), bears a striking resemblance to the phenomenon of liquid phase condensation^88^ as a means to form P-bodies, stress granules, and other types of intracellular protein gels or “membrane-less organelles”. In such liquid-liquid phase separated systems^89^, a multivalent web of relatively weak (individually) molecular interactions leads to the mesoscopic assembly of a distinct, demixed liquid phase (e.g., the nucleolus) within the cell. Notably, these molecular interactions generally occur between low-complexity, conformationally pliable peptides, as in the recently characterized, hydrogel-forming “low-complexity aromatic-rich kinked segments (LARKS)”^90^. A possible direction for future work involves elucidating any similarities between the ‘aromatic ladders’ and other structural features of LARKS assemblies and, for instance, the conserved Phe in our RAPID peptide systems.

### Electron Microscopy of Nanofiber Morphology: TEM and CryoEM

Fibrils, tubes, dendrimers and other ultrastructures often form via a hierarchical supramolecular arrangement of specific, non-covalent contacts^91–92^. TEM analysis revealed that our RAPID hydrogels are composed of nanofibers as well as dense regions of fibrous bundles. At low pH (i.e. non-gelling conditions), fibers do not form within the peptide solution; rather, amorphous aggregates are present (Figure 7A). At physiological pH, individual fibers bundle into hierarchical nanostructures with clearly twisted, ribbon-like morphologies (Figure 7B). The multi-stranded, twisted ribbons reported here are unique among nanofiber-forming, self-assembling peptide hydrogels^41, 93^. In at least some characterized systems, the helicity (and other geometric properties) of fibers are thought to depend on such atomic-level effects as the properties of steric packing between aromatic side-chains, such as for Tyr and Phe^94^; whether the general morphological properties that we find for RAPID peptides can be traced to such underlying factors is an appealing question for future structural modeling studies. In earlier work^87, 94^, cooperative intermolecular hydrogen-bonding between the backbone N- and C-termini were found (by modeling) to enable stronger interactions (i.e. closer intermolecular packing), leading to the classical geometric features of twisted ribbons. Our peptides are C-terminally amidated, and it is more likely that RAPID fibrils assemble via anti-parallel stacking of pentapeptides, with details of the molecular packing predominantly stemming from apolar dispersion forces and other enthalpically favorable interactions among the Phe moiety and amphiphilic nature of the sequence^95–96^.

**Figure 7.**
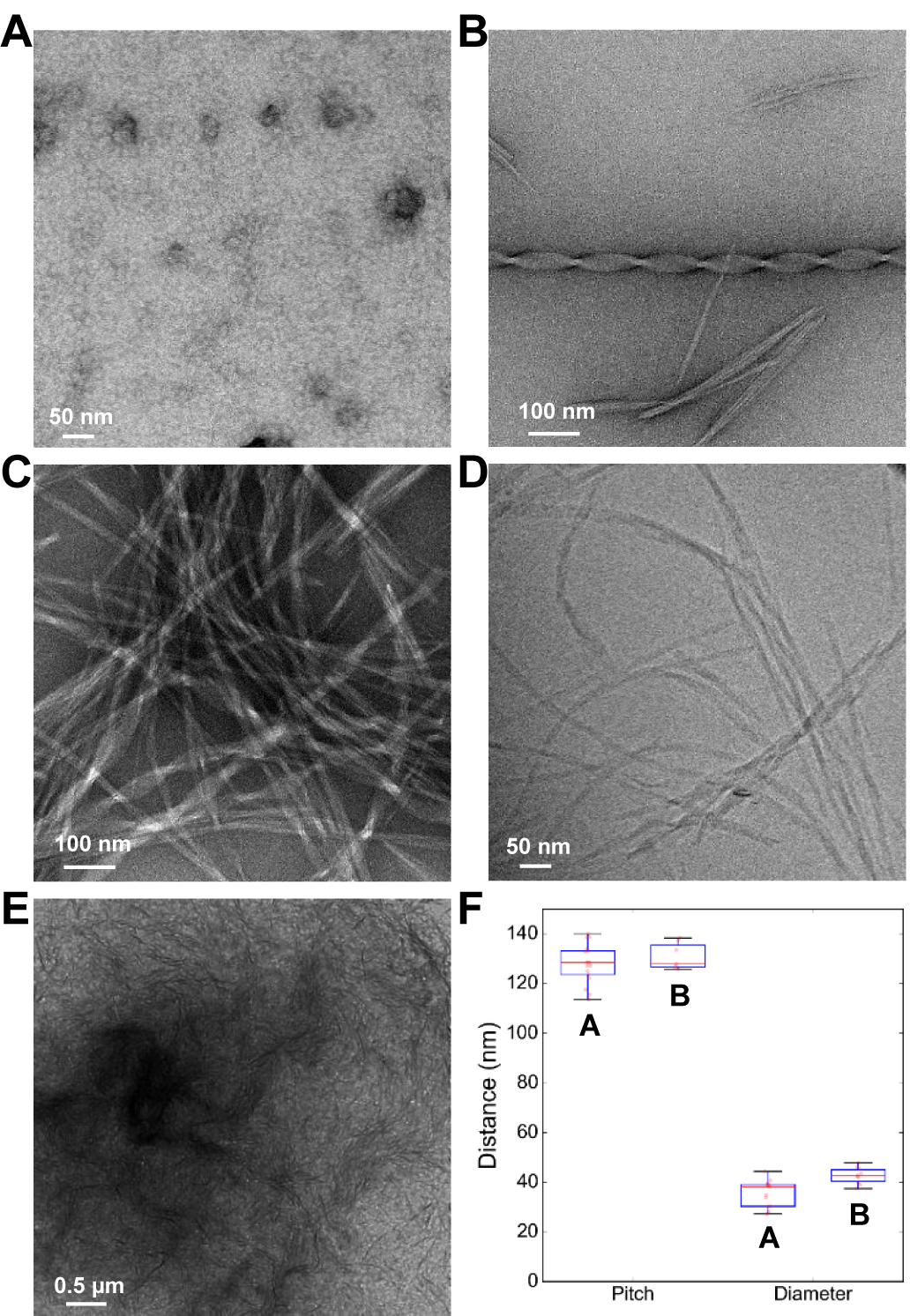
Representative EM images of 1.5 wt.% KYFIL hydrogels. A) Images of amorphous peptide aggregates in non-gelling conditions (pH 4.6). There is no distinct fiber formation within peptide solutions. B) Images of individual twisted ribbon molecular assemblies present within the hydrogel at pH 7.4. These twisted ribbons have ca. 40 nm width and ca. 132 nm pitch. C) TEM images of bulk fibres within the hydrogel. Both ‘classical’ fibrous bundles that are commonly observed in other reported self-assembling peptides and the twisted ribbon morphology are present within this hydrogel system. D) Cryo-EM images of 1.5 wt. % KYFIL hydrogel. Twisted ribbon morphologies are present within the hydrogel. E) Lower magnification of the KYFIL peptide, demonstrating that twisted ribbon morphologies are present in mass throughout the hydrogel volume. F) Quantification of the pitch and diameter of the twisted ribbons is consistent and reproducible. A and B refer to different synthetic batches.

Individual fibers can apparently entangle, yielding multistranded twisted ribbons (Figure 7C). Similar hierarchical ‘bundling’ of fibrils, interwound ‘superhelices’, and other higher-order assemblies have been seen in systems such as amyloid-related peptides^97–99^. In other previously characterized self-assembling peptide systems, ionic interactions, modulation by the solvent environment, and hydrogen bonds are thought to govern the formation of interconnected networks of nano-fi-brils^13, 15, 24, 27, 93, 100–101^. In addition—unlike other nanofiber-forming peptides—the pentapeptide sequences presented here are significantly shorter than many hitherto characterized systems (decapeptides and beyond). Furthermore, other self-assembling peptide hydrogels^13, 24, 85, 92^ often lack distinct morphology within their nanofiber-forming sequences (instead being irregular and heterogeneous), whereas RAPID peptides form highly regular, twisted fibril nanostructures.

Cryo-TEM of our pentapeptide samples in vitreous ice reveals fibers that maintain twisted fiber morphologies, with finite fiber lengths up to ∼100 μm (Figure 7D). At relatively low magnification, twisted ribbons appear to pervade the hydrogel network, suggesting that these particular morphologies are not isolated, localized or otherwise spurious instances of self-assembly (Figure 7E). As part of an unbiased experimental design, two different batches of the KYFIL pentapeptide were independently synthesized and evaluated under TEM in order to assess the robustness and reproducibility of fibril formation. The geometric features of the fibers (helical pitch, diameter) were consistent across the two separate batches (Figure 7F), demonstrating that these results are replicable and that there is low batch-to-batch variability as regards the peptide synthesis process, purification, and self-assembly. The periodicity of the fibrillar twist is ≈ 120 nm, as determined via visual analysis (Figure 7) and by calculation of the autocorrelation function of pixel intensity along individual fibrils (Figure S12).

Notably, both of these morphological features of our RAPID peptides—the presence of a fibrillar twist, and the ≈ 120 nm value of its pitch—are recapitulated in the structural features of many peptide-related systems, amyloidogenic and otherwise. One example is β-sheet fibrils and “periodically-twisted nanoribbons” formed by an Ac-NNFGAILSS peptide from the “amyloidogenic core” of islet amyloid polypeptide (IAPP_21-29_)^97^. These peptides featured axial repeats of ≈ 85-100 nm. In a closely related system, an overlapping IAPP-derived peptide (IAPP_20-29_) had AFM-characterized fibril periodicities of ≈ 203 nm^99^. Fibrils from disparate proteins (e.g., SH3-containing proteins, and lysozyme) can also be polymorphic. Based on AFM studies, two subpopulations of SH3 fibrils form helical repeats of ≈ 105 nm and ≈ 156 nm, while human lysozyme fibrils have an “axial crossover repeat” of ≈ 200 nm^102^. Perhaps most pertinent to our current study, systematic studies of a family of short peptides based on I_3_K (including all stereoisomeric combinations of L- and D-amino acids), showed that these amphipathic peptides form twisted fibrils with a helical pitch of ≈ 120 nm^103^. This is in remarkable agreement with our RAPID fibrils, which exhibit nearly the same pitch (Figure 7F). Though not identical to these previously characterized systems, the morphological properties of our RAPID peptide-based fibrils nevertheless are quite similar, suggesting that perhaps some unifying structural and energetic principles underlie the formation of these various supramolecular structures, amyloid-related and beyond. Most broadly, such commonalities could have overarching implications for peptide engineering and nanomaterials.

As expected, peptide sequence has a significant effect on the nanofiber morphology. More specifically, the self-assembly of hierarchical twisted ‘macromolecular’ structures can be altered by substituting any residue within the …Phe–Ile–Leu… moiety that detracts from the amphiphilicity of the sequence and π-system interactions. Similarly, any modification to the sequence also results in drastically different mechanical properties, as indicated in our rheology studies. We observe some twisting in nanofibers occurs within 1.5 wt. % AYFIL hydrogels at pH 7.4, but the typical diameters of these fibers (≈ 10 nm) are significantly smaller than those of KYFIL hydrogels (≈ 40 nm). Though impossible to assess without more detailed analyses, a possible molecular basis for this difference relates to the sterically smaller alanine enabling a tighter packing of individual peptides within fibers or protofibers (versus the more extended Lys side-chain). For the KAFIL system under the same conditions, there is no distinct fiber formation—only spherical aggregates are seen (Figure 8), though it should be noted that KAFIL peptides can form hydrogels at higher pH conditions. Interestingly, for KYFAL hydrogels, twisted ribbon morphologies still occur, though the persistence length of these fibers appears to be significantly shorter (based on qualitative/visual analysis), and the twist periodicity (i.e., helical pitch) is more irregular. The change in fibrillar morphology, upon an Ile → Ala substitution, may ultimately stem from an alteration in the steric properties of side chain–mediated geometric packing of peptides. While there is a great difference in length-scale between an individual peptide on the nm-scale and a supramolecular assembly (such as a fibril), we do see correlations between hydrophobicity properties of the different pentapeptides and the patterns of relative solvent accessibility across the different peptides, as captured by MD simulations (Figure 5). An intriguing problem for future work is elucidation of the sequence correlates and stereochemical basis for fiber morphology (e.g., thicker ribbon diameters [≈40 nm] for KYFIL versus [≈10 nm] for KYFAL). Successfully addressing this goal will likely require an integrative, multidisciplinary and multiscale approach, such as was used to decipher the atomic structures of cross-β amyloid fibrils of a transthyretin-derived peptide^98^.

**Figure 8.**
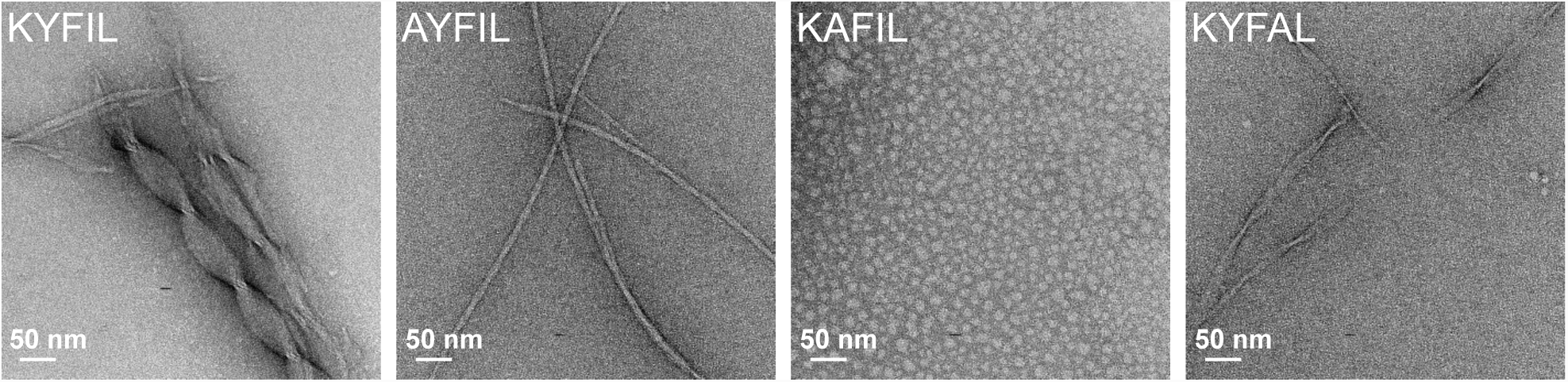
Representative TEM images of 1.5 wt. % pentapeptides in PBS at pH 7.4. KYFIL hydrogels exhibit twisted ribbon morphologies, while AYFIL hydrogels are comprised of twisted fibrils. KAFIL peptide solutions at pH 7.4 form spherical aggregates (non-gelling conditions), while KYFAL hydrogels also form twisted ribbon morphologies, with longer and more infrequent pitch than KYFIL peptides.

### Cell-Protection by RAPID Hydrogels during Syringe Ejection

During syringe needle flow, cells can experience various types of mechanical forces that ultimately disrupt the cellular membrane: 1) extensional flow, where cells encounter stretching forces, 2) pressure drop across the cell, and 3) shear stresses, due to linear shear flow as the cell travels across the syringe^6^. In our present study, we experimentally tested the effects of syringe needle flow on the viability of oligodendrocyte precursor cells (OPCs) suspended in PBS or RAPID hydrogels as a cell carrier at a flow rate of 1000 µL/min. OPCs are therapeutically relevant, as OPC transplantation may help circumvent the inherent regenerative limitations within the central nervous system (CNS).^104–105^ Indeed, this OPC transplantation strategy is currently being pursued as a therapeutic intervention in human traumatic spinal cord injury patients^106^. When cells were ejected in PBS, OPC viability was significantly decreased compared to cells encapsulated in RAPID hydrogels and ejected (p < 0.05, Figure 9). This finding suggests that RAPID hydrogels could protect transplanted cells from the mechanical forces encountered during syringe needle flow and serve as valuable cell carriers in transplantation protocols.

**Figure 9.**
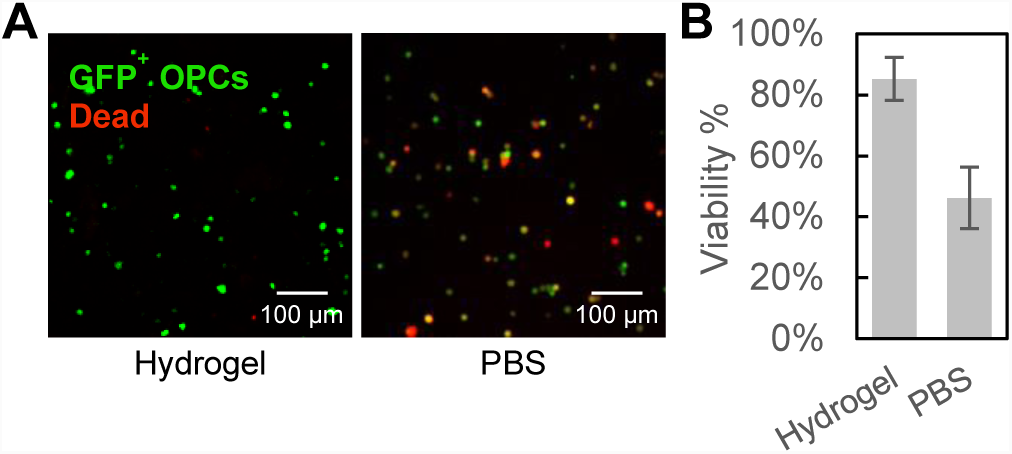
Viability of OPCs immediately after syringe needle flow in PBS and AYFIL hydrogels. A) Live/Dead images of viable (green, GFP+) and membrane damaged (red, ethidium homodimer-1) cells postejection in PBS or 1.5 wt. % AYFIL hydrogels. Each sample of cells encapsulated in RAPID and PBS, respectively contained at least 140 total cells. B) Percent cell viability with injection in PBS or hydrogels. Error bars represent standard error of the mean (SEM) from three separate syringe ejections (n = 3), *p < 0.05.

### Cytocompatibility and 3D Cell Culture Potential of RAPID Hydrogels

Mounting evidence now highlights the mechanosensitive nature of OPCs within the CNS. These lineage-restricted glial cells give rise to myelinating oligodendrocytes, and OPC proliferation and differentiation both correlate with the physical stiffness of underlying 2D^107^ or surrounding 3D matrices^108^.

The biophysical properties of hydrogels sharply influence the proliferation and differentiation of stem cells within a 3D environment^109–110^. For instance, neural stem cells (NSCs) proliferate significantly more in softer substrates^1, 111^, and preferentially differentiate into neurons in hydrogels with low moduli^112–114^. Recent evidence indicates that OPCs are also sensitive to the biophysical stiffness of their surrounding microenvironment^115^. We encapsulated OPCs^116^ in order to examine the effect of the RAPID hydrogels on viability and proliferation. OPCs survived and grew in 1.5 wt. % AYFIL hydrogels (1900 Pa), as determined by the increase in both ATP and DNA concentrations over time (Figure 10a and 10b, respectively). A 1.5 wt. % AYFIL hydrogel was used to investigate cytocompatibility and cell growth, as its mechanical properties (∼1900 Pa) approximate CNS tissue stiffness^1, 117^. Cell encapsulations with 1.5 wt. % KYFIL hydrogels resulted in poor cell viability, likely due to the stiffness (∼8000 Pa) being much greater than native CNS tissue.

**Figure 10.**
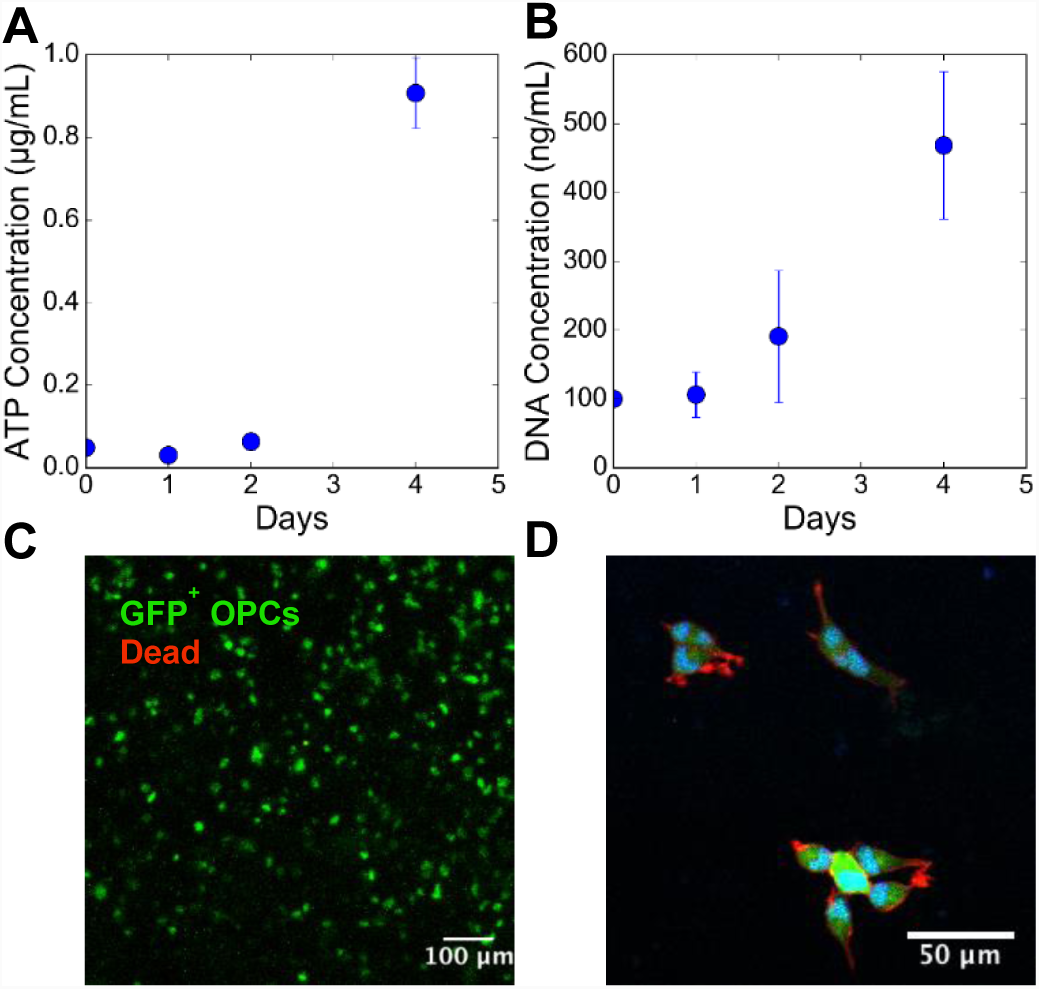
MADM OPC line encapsulated in 1.5 wt. % AYFIL hydrogels and cultured over 4 days. A) OPCs remained viable after encapsulation for at least 4 days, as determined by the increase of ATP over time (B). The increase in DNA concentration suggests that cells proliferate over the course of 4 days. Error bars represent standard error of the mean (SEM, n = 3). C) Live/Dead (green/red staining) images taken at Day 1 of the experiment demonstrate that the majority of cells remained viable following encapsulation. Image is a maximum projection of a 132 µm thick z-stack. D) Maximum projection (23 µm thick z-stack) of OPCs encapsulated in AYFIL hydrogels after 2 days of culture. Process extension of OPCs are observed, suggesting that these hydrogel systems are suitable for neural cell culture. GFP (green), Actin (red), DAPI (blue).

Live/dead imaging indicated a high percentage of viable cells (Figure 10C). Others have previously shown that OPCs can extend processes within 3D poly(ethylene glycol) hydrogels after 7 days of culture, but only in the presence of laminin^108^. Here, we demonstrate that cells within an AYFIL hydrogel can extend processes within 2 days of culture without any bioactive cellular adhesion peptide sequences or incorporating native ECM proteins (Figure 10D, Movie S8-S10). This could be due to physical hydrogel peptide matrix being permissive of remodeling by the cells. This finding highlights that the simplicity of our cell culture system is sufficient for growth of cells derived from the CNS, without the need for laminin-derived peptide sequences as has been demonstrated in other peptide hydrogel systems^15^.

## Conclusions

We have devised a new family of short, five amino acid, peptide sequences capable of self-assembling into robust hydrogels. We synthesized seven closely-related, stimuli-responsive pentapeptide sequences. Four of our RAPID sequences form robust hydrogels at concentrations down to at least 1.5 % (w/v). Physicochemical features of the sequence—in particular, amphiphilicity and inclusion of a central phenylalanine—influence the self-assembly and β-strand formation propensities of this class of peptides. MD simulations, aimed at examining the structural properties of these β-strand–forming peptides, reveal that hydration plays an integral role in the conformational dynamics of these peptides. Experiments reveal that our hydrogels exhibit shear-thinning and self-healing properties—features that may stem, at least partly, from the facile formation of β-sheet structures (in accord with our MD simulations). These rheological properties suggest the suitability of our RAPID peptides for biomedical applications requiring injection. Additionally, we observe that at physiological pH, hierarchical nanostructures (i.e. individual fibers) bundle into clearly twisted, ribbon-like morphologies. The multi-stranded, twisted ribbons reported here are unique among nanofiber-forming, self-assembling peptide hydrogels. We demonstrate that these self-assembling hydrogels offer effective strategies for encapsulating OPCs within 3D matrices of tunable viscoelasticity. These scaffolds allow for cell growth and morphological process extension in OPCs. We also demonstrate that RAPID hydrogels can mitigate the damaging effects of extensional flow during syringe injections. The supramolecular assemblies formed by RAPID peptides represent injectable hydrogel systems that may offer new and translational approaches for cell delivery and tissue engineering applications.

## ASSOCIATED CONTENT

### SUPPORTING INFORMATION

Materials, equipment, synthesis and hydrogel characterization, MD simulation system setup, and Figures S1–S19. Table of MD simulation systems, representative snapshots of simulated systems, radius of gyration, Ramachandran plots, other rheological characterization, periodicity of fibrillar twist as quantified by intensity auto-correlation function, and MALDI-TOF spectra of all synthesized pentapeptides, trajectory movies, cell projection Z-stack movies, hydrogel forming movie, hydrogel syringe ejection movie.

## AUTHOR INFORMATION

The manuscript was written through contributions of all authors.

All authors have given approval to the final version of the manuscript.

## Notes

The authors declare no competing financial interest.

## ACKNOWLEDGEMENT

The authors thank UVA’s Molecular Electron Microscopy Core for EM imaging and UVA’s W.M. Keck Center for Cellular Imaging for use of the Zeiss 510 and Zeiss 780 confocal/multiphoton microscopy systems. We acknowledge support from the UVA Center for Advanced Biomanufacturing for acquiring and supporting the peptide synthesis and rheometry equipment. We also thank Dr. Phil Bourne (UVA) for supporting these efforts, as well as Stephanie Guthrie and Dr. Gaurav Giri (UVA) for their assistance with FTIR characterization.

## Materials & Methods

### Peptide Synthesis

All peptides were synthesized by solid-phase chemistry in 0.1 mmol batches on a Tribute peptide synthesizer (Gyros Protein Technologies, AZ). A TentaGel R Rink Amide Resin was used which results in a C-terminal amide. Solvents and Fmoc (fluorenylmethoxycarbonyl)–protected AAs were purchased from Gyros Protein Technologies. Reagents were made with 5 equiv. moles of amino acid and 5 equiv. moles of HBTU (2-(1*H*- benzotriazol-1-yl)-1,1,3,3-tetramethyluronium hexafluorophosphate), and subsequently dissolved in DMF (dimethylformamide). Amino acid coupling cycles were 60 min in length. Protecting groups were removed with treatments of 20/80 v/v piperidine/DMF for 10 minutes. After the coupling reaction was complete, the resin was washed three times with DCM (dichloromethane) before running the cleavage step. Cleavage of the peptides were accomplished by shaking the resin with 10 mL of TFA (trifluoroacetic acid)/triisopropylsilane/H_2_O (95:2.5:2.5 volume ratios) for 2 h at room temperature. The peptide solution was collected, and the peptide precipitated by the addition of cold diethyl ether followed by two washes with cold ether after centrifugation. The peptides were dried overnight, redissolved in deionized water and dialyzed with 10 water exchanges over 5 days using molecular weight cutoff of 100-500 Da (Spectra/Por, Spectrum Laboratories Inc., Rancho Dominguez, CA). Lyophilized peptides were stored at −20 °C and protected from light. MALDI-TOF (matrix assisted laser desorption ionization time-of-flight) analysis was used to characterize the mass of the final products. See Supporting Information for spectra (Figure S7-S13).

### Attenuated Total Reflection–Fourier Transform Infrared Spectroscopy (ATR-FTIR)

IR measurements were obtained for 3 wt. % peptides in PBS on a PerkinElmer 400 FT-IR spectrometer equipped with an ATR accessory. Aliquots of the peptide were deposited on a “Golden Gate” diamond ATR (PerkinElmer, USA). PBS was used as a background spectrum. Collected spectra were normalized by dividing all the absorbance values in the spectrum within the Amide I band by the largest absorbance value^1^, baseline corrected, and vertically offset for ease of comparison.

### Molecular Dynamics Simulations of Pentapeptides

KYFIL, KYFAL, KAFIL and KYFIV peptides were constructed using the peptide builder tool in the program *Avogadro*^*^2^*^. A custom Tcl script was used to amidate the C-termini in VMD^3^ using the CHARMM36 forcefield^4^. Eighteen individual pentapeptides were solvated in a cube of explicit TIP3P water molecules, using VMD’s solvation box extension; a 4-Å padding between the solute and nearest box face was used along with periodic boundary conditions. The pentapeptides were staggered 8 Å apart (as measured by their geometric centers) and rotated randomly so as to prevent orientational bias in the starting structures. The final simulation cell contains approximately 15,000 atoms (and varies with peptide sequences) with a rectangular parallelepiped box of water measuring 67 Å × 71 Å × 49 Å. Physiological concentrations (150 mM) of Na^+^ ions, including sufficient Cl^−^ ions to neutralize the solute’s charge, were added to the solvated system using VMD’s *ionize* plugin. The internal energy was minimized for 10,000 steps, and the system was then equilibrated for 10 ns (with a 2-fs integration step) in the isothermal–isobaric ensemble (NPT) ensemble. Temperature (300 K) and pressure (1 atm) were regulated via Langevin dynamics for all non-hydrogen atoms and a hybrid Nosé–Hoover Langevin piston. Simulations were performed in NAMD 2.10^5^, with final production trajectories extended to 200 ns. Trajectories were processed and further analyzed using in-house scripts written in the Python^6^ and D^7^ programming languages, as well as VMD. Secondary structures were assigned using STRIDE^8–9^. Peptide structures were characterized via the SURF calculation (surface areas), with the solvent probe radius set to 1.4-Å and applied to all peptides; in this way, clusters were then defined as any peptides that have overlapping molecular surfaces, within 1.4-Å of each other. Density function plots were determined using a univariate kernel density estimate from the Python Seaborn package. Table S1 summarizes our peptide MD simulation systems. Grand average hydropathicity (GRAVY) values are computed from Kyte-Doolittle (KD) hydrophobicity^10^ indices averaged over the amino acid sequence for each peptide.

### Hydrogel Formation and Rheological Properties

Lyophilized peptides were dissolved in PBS to a final concentration of 1.5 or 3 wt. %. The pH of the peptide solutions was adjusted by drop-wise addition of minute amounts of HCl or NaOH. Rheological tests were performed on 50 μL hydrogels 30 minutes after induction of gelation (Anton Par, P25S 25 mm parallel steel plates) with a measuring gap of 250 μm. Storage (*G*′) and loss (*G*”) moduli were measured as a function of strain (%) ranging from 0.01 to 100% with a constant frequency of 10 rad/s. Frequency sweeps were performed at angular frequencies ranging from 1 to 100 rad/s at 0.1% strain. For recovery experiments, a step-time procedure was utilized with a series of applied strains at a fixed oscillation frequency of 10 rad/s. Initially, samples were applied with 0.01% strain for 100 s followed immediately by a 100% strain for 50 s, and cycled 5 times.

### Transmission Electron Microscopy

3.5 μL of peptide hydrogel was placed on a holey carbon grid (Protochips, Inc.). Three washes of deionized water, and three washes of 2% uranyl acetate staining solutions with 2 s blotting between each step was performed. Samples were analyzed on a Tecnai F20 equipped with a 4k × 4k UltraScan CCD camera. Pitch and length were determined using Fiji^11^, and the FiberApp software package^12^ was used to compute autocorrelation functions of intensity profiles.

### Cell Culture

GFP^+^ MADM OPC lines^13^ were expanded *in vitro* on T75 tissue culture plates treated with poly-ornithine. OPCs were cultured in DMEM with high glucose, 4 mM L-glutamine, 1 mM sodium pyruvate (Life Technologies) with N2 and B27 supplement (Life Technologies), 1% penicillin-streptomycin (Life Technologies), 10 ng/mL mouse PDGFA-AA (eBioscience), and 50 ng/mL human NT3 (Peprotech). Cell media was changed every 2 days, and cells were grown to 90% confluency and passaged using 0.25% trypsin in Dulbecco’s phosphate buffered saline (PBS). Cells were cultured in 5% CO_2_ atmosphere, and 21% O_2_ at 37 °C.

### Hydrogel Cell Encapsulation and Analysis

Hydrogels for cell encapsulation were made using 1.5 wt% AYFIL peptide in PBS. 25 uL hydrogels with 5 × 10^6^ cells/mL were made by mixing cells and peptides and then transferred to a cell incubator for 10 minutes at 37 °C at 100% humidity. OPC proliferation media was then added to the hydrogels, and changed every 2 days. Hydrogels were stored at −80 °C in lysis buffer before running ATP or DNA quantification assays. For quantification, gels were homogenized in lysis buffer using a hand grinder and were measured using the CellTiter-Glo luminescent Cell Viability Assay (Promega, United States) and the Quant-iT PicoGreen dsDNA assay (ThermoFisher) according to manufacturer protocols.

### Syringe Needle Flow Viability Assay

OPCs were resuspended at a cell density of 1 × 10^5^ cells/mL in either PBS or 1.5 wt.% AYFIL hydrogels, and loaded into a 1-mL syringe with an 18-gauge needle, mounted onto the syringe pump, and ejected onto a 24 well plate at a constant volumetric flow rate of 1000 µL/min. Cell viability was assessed with a dead stain assay (Invitrogen). Briefly, hydrogels were rinsed for 10 minutes in PBS plus glucose (PBSG), and stained with 4 μM ethidium homodimer-1 for 40 minutes in PBSG, and rinsed in PBSG prior to imaging. GFP^+^/dead images were collected using a Zeiss LSM 510 confocal microscope. 150 μm z-stack images were collected with a frame distance of 1 μm. For image analysis, channels for live cells in the green channel and dead cells in the red channel were split and analyzed separately, and converted to 8-bit to allow for thresholding based on the intensity. The Find Maxima plugin was used for each channel to count the number of dead or live cells. Using the point selection tool, the Noise Tolerance values were adjusted by increments of 5 until background staining was excluded (the Noise Tolerance value was 50 for all images). The number of points for each channel were recorded, and were used to calculate the percentage of live and dead cells. Total cell count was between 146 and 287 for each image (n = 3 samples per condition). Statistical significance was determined using the Student’s t-test with p < 0.05.

### Immunostaining

After 2 days of culture, gels were fixed in 4% paraformaldehyde for 20 minutes at 4 °C and rinsed with PBS before permeabilizing overnight with 0.3% triton-X in PBS. Hydrogels were rinsed in PBS, and then incubated with 10 µg/mL stock solution of Alexa Fluor 568 Phalloidin (ThermoFisher) in 1% BSA in PBS overnight. 4′,6-diamidino-2-phenylindole (DAPI) was added to stain cell nuclei during the last 20 minutes of incubation. Gels were then washed 4 × 20 min in PBS and imaged with a Zeiss LSM 780 confocal microscope. 100 μm z-stack images were collected with a z-spacing distance of 1 μm.

---

**Table 1.**
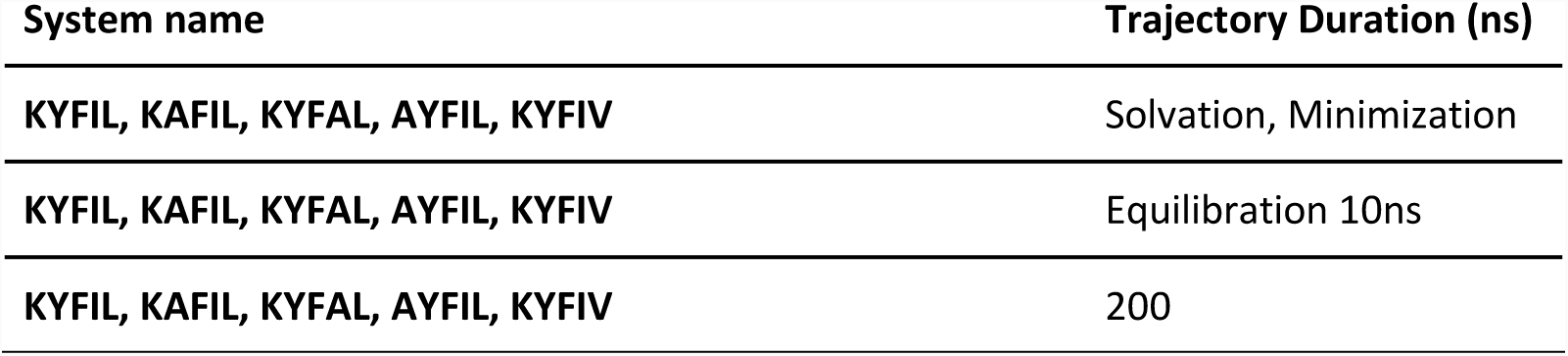
Summary of MD simulation systems of the pentapeptide systems

**Figure S1.**
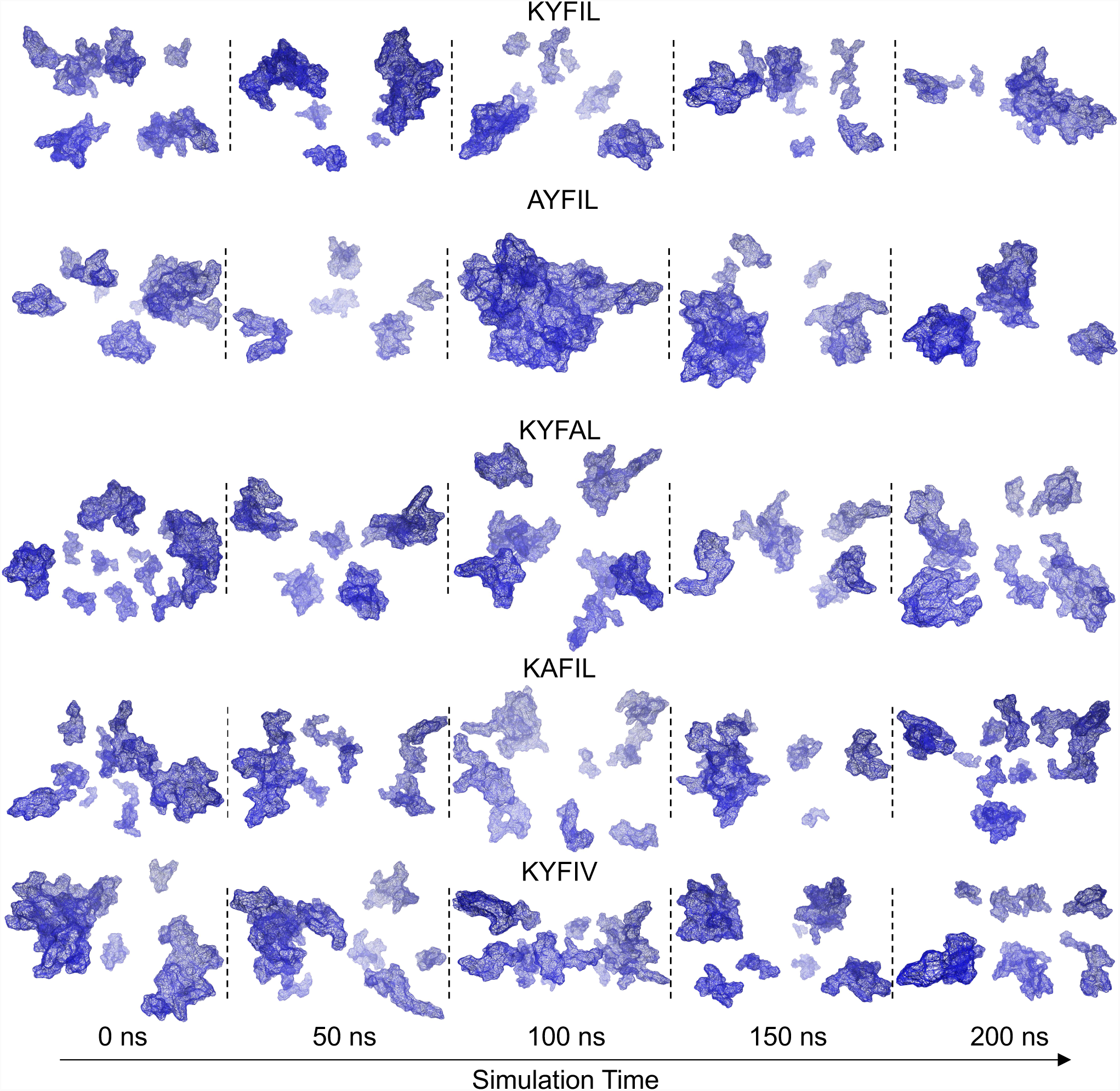
Representative snapshots of KYFIL, KYFAL, KAFIL, and KYFIV peptide systems at increasing time points following minimization and equilibration. Snapshots were taken after minimization for 10,000 steps, and equilibration for 10 ns. MD simulations were conducted for 200 ns, and peptide systems were simulated with an explicit water solvent (TIP3 solvent model). For experimentally-determined gelling peptides (KYFIL, AYFIL, KYFAL, KAFIL) the number of peptide clusters decreases as the simulation progresses, highlighting their aggregation propensity.

**Figure S2.**
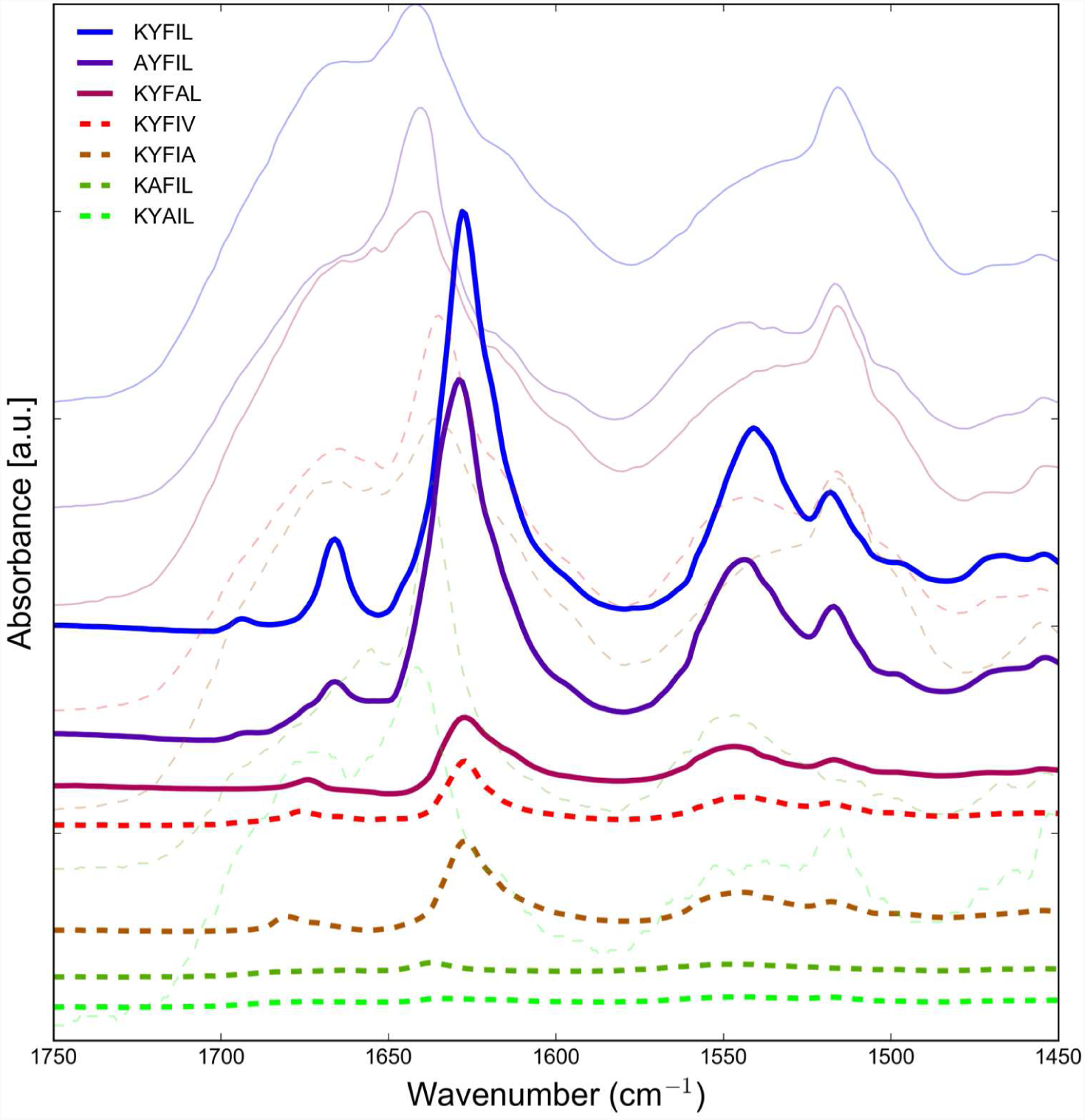
ATR-FTIR spectrum of peptides in PBS (thick lines) and freeze-dried peptides (thin, transparent lines). All peptides that are able to gel at pH 7.4 (solid lines) exhibit an Amide I absorbance at 1629-1645 cm^−1^, indicative of β-sheet hydrogen bonding. Non-gelling peptides in the same conditions (dashed lines) exhibit much weaker, less intense peaks at the same wavenumbers. All spectra are baseline corrected, normalized, and offset for clarity.

**Figure S3.**
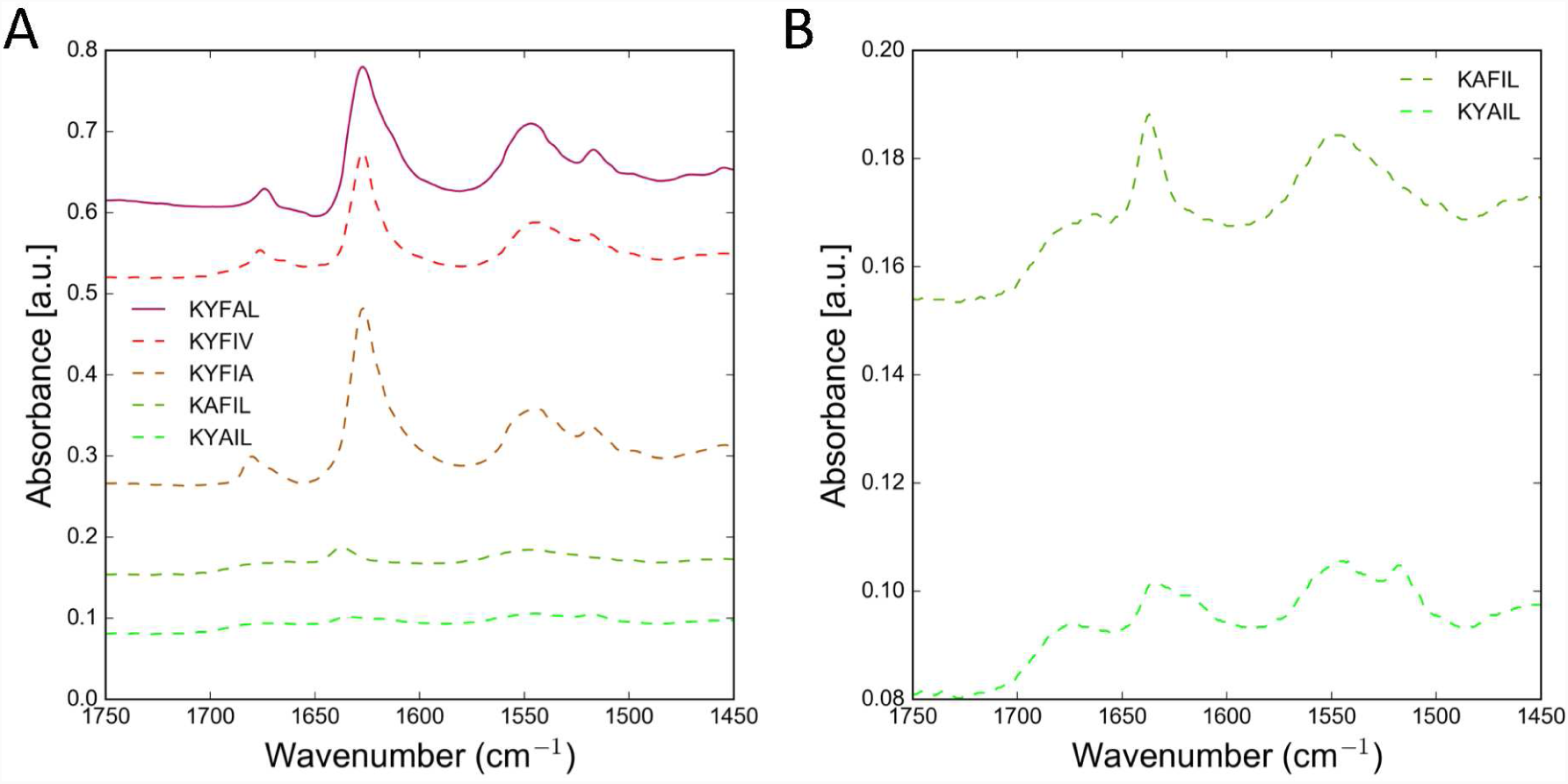
A) A magnified view of the ATR-FTIR spectrum of peptides dissolved at 3 wt.% in PBS and pH 7.4. All gelling peptides (solid lines) exhibit an Amide I absorbance at 1629 cm^−1^, indicative of β-sheet hydrogen bonding. A peak at 1679 cm^−1^ to 1683 cm^−1^ indicates that the -sheet is in anti-parallel conformation. B) Non-gelling peptides (dashed lines) exhibit much weaker, less intense peaks at the same wavenumbers. All spectra are baseline corrected, normalized, and offset for clarity.

**Figure S4.**
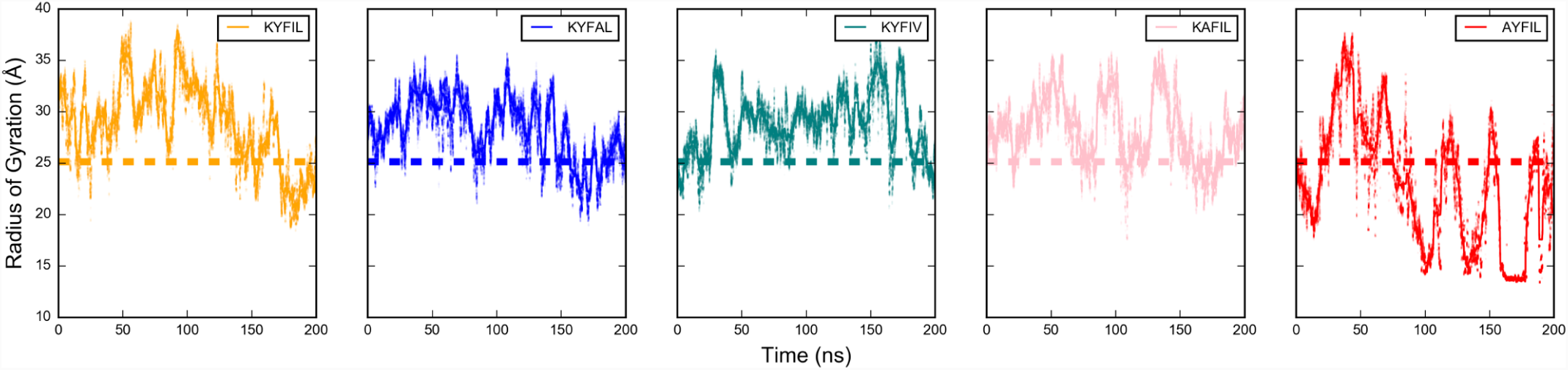
Sequence dependence of the radius of gyration (R_g_). The R_g_ was measured for an ensemble of 18 peptides of different sequences (KYFIL, KYFAL, KYFIV, KAFIL, and AYFIL) after equilibration of 10 ns. All peptides incur hydrophobically-driven collapse (relative to initial starting structure). Dashed line indicates initial R_g_ before equilibration. Towards the end of the simulation, the R_g_ for KYFIL and AYFIL decreases relative to the beginning of the trajectory, highlighting their aggregation propensity.

**Figure S5.**
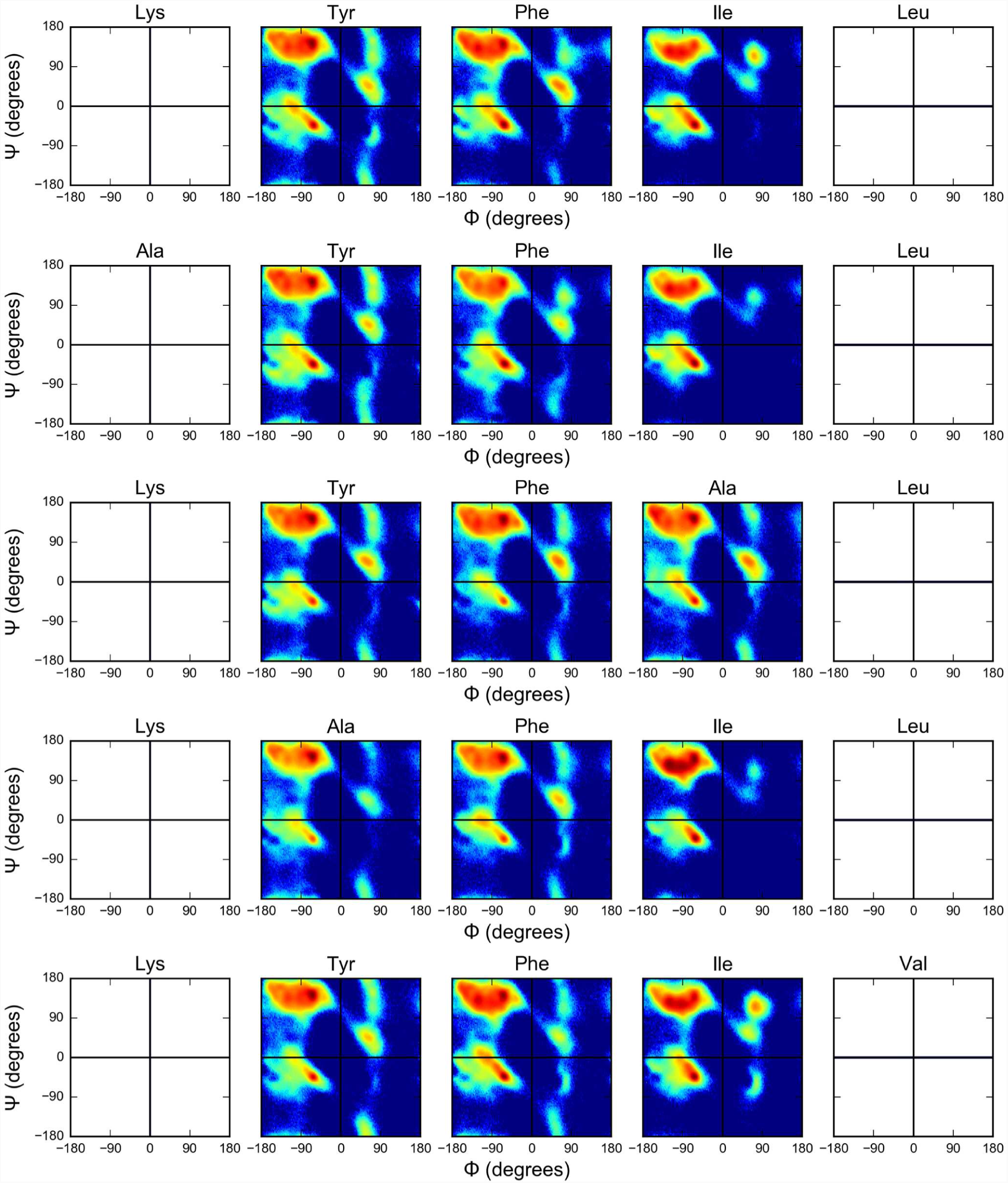
Ramachandran plot (ϕ, ψ distributions) for each residue in a pentapeptide sequence. The torsion angles for each type of amino acid, barring the N- and C-termini indicate significant structural heterogeneity within the peptide systems. The Phe for all pentapeptide analogs, adopts higher populations of *B*-turn type-II(ϕ= −60 °, φ= 120°) and antiparallel *B*-sheet structures(ϕ= − 140°, φ = 135 °). For both the KYFAL and KAFIL sequence, the Ala preferentially adopts a polyproline type-II helix(ϕ= −75 °, φ = 145 °) and decreased propensity for *B*-sheet structures. Note that the residue at the N- and (-terminus does not have a Phi or Psi angle since the dihedral angle requires a plane comprised of C′-N-Cα-C′ and N-Cα-C′-N for Phi and Psi angles, respectively.

**Figure S6.**
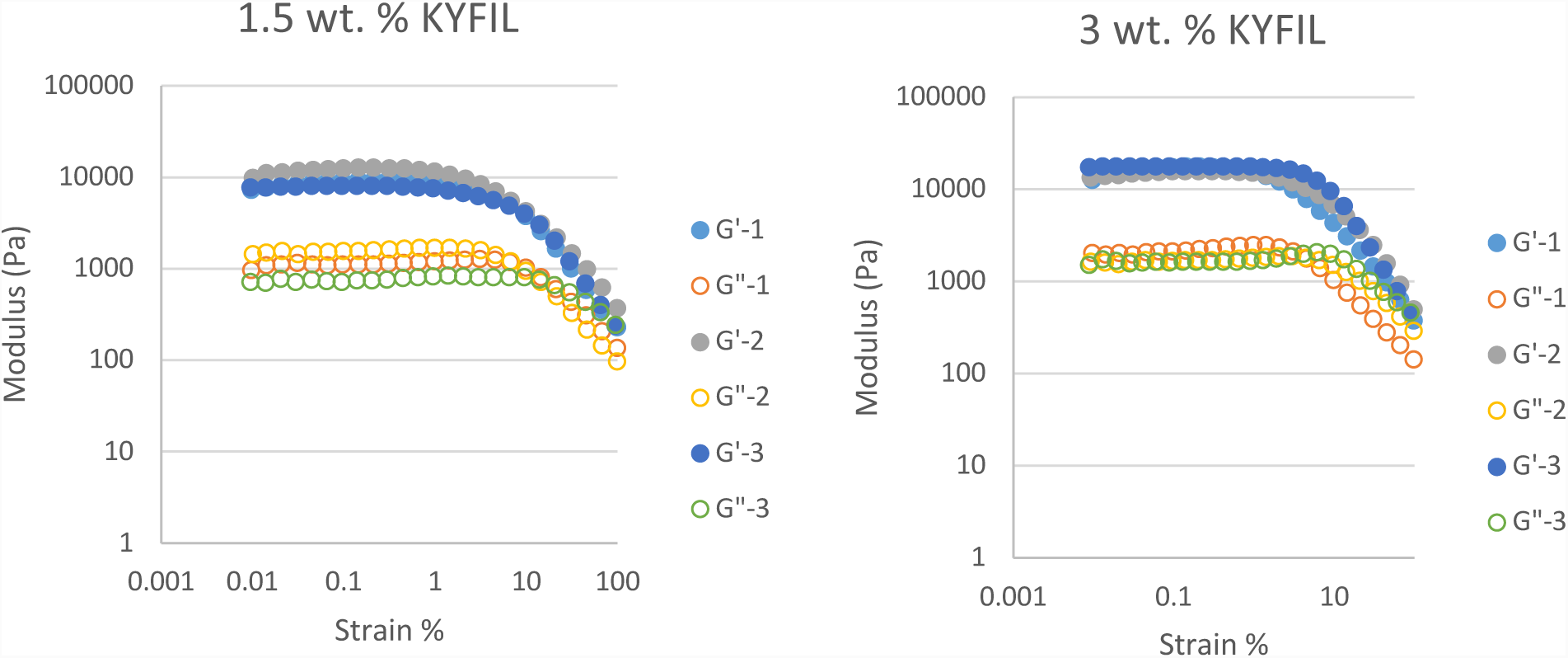
Strain sweep of gelling KYFIL sequences at constant frequency of 1 hz. Measurements are carried out at 3 wt.% and 1.5 wt %. For all sequences, G′ decreases significantly in acidic conditions. Higher concentrations of peptides exhibit increased G′. Legend indicates 3 sample replicates.

**Figure S7.**
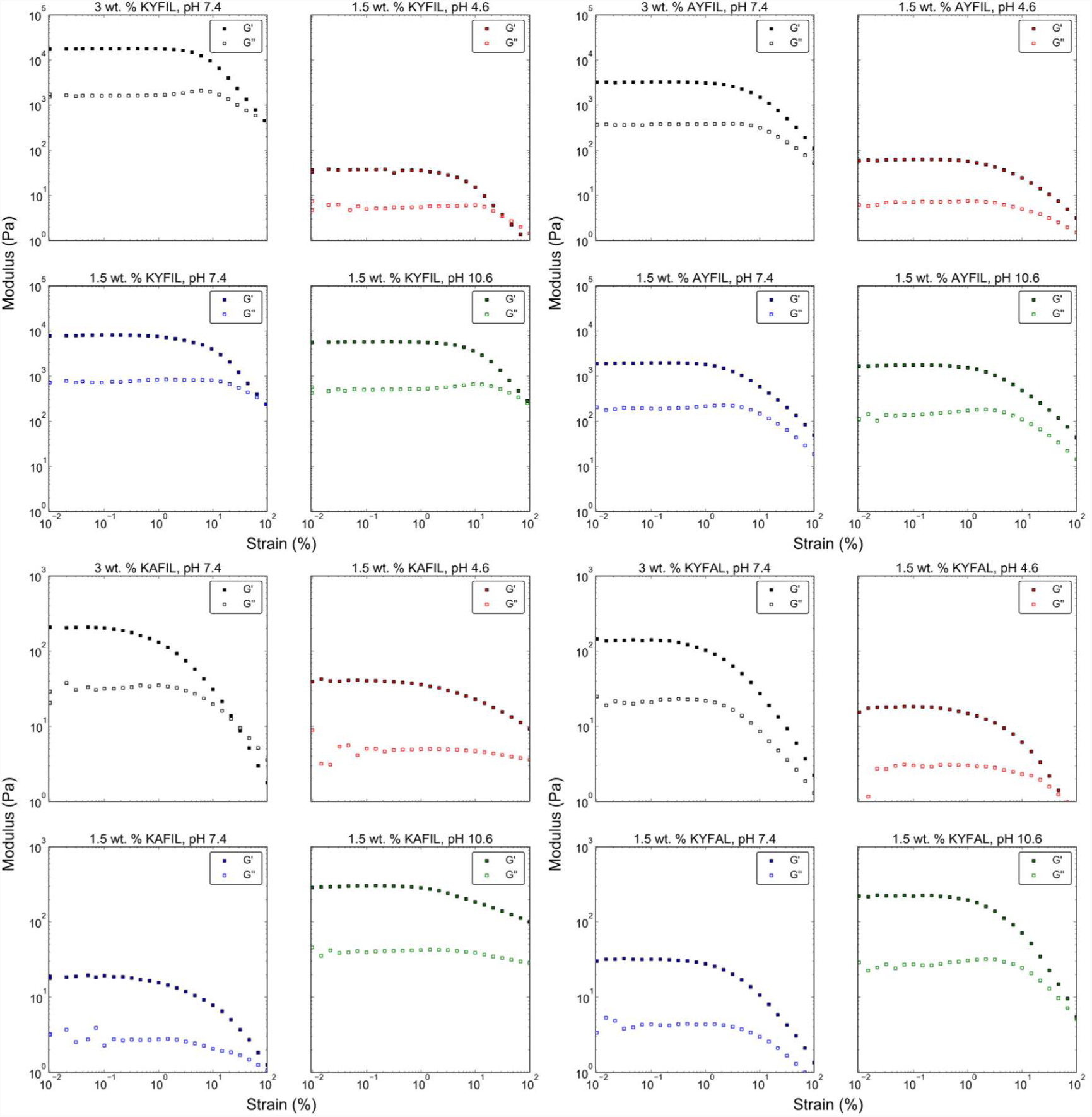
Strain sweep of gelling pentapeptide sequences at constant frequency of 1 hz. Measurements are carried out at 3 wt.% and 1.5 wt.%, and different pH conditions (4.6, 7.4, 10.6). For all sequences, G′ decreases significantly in acidic conditions. Higher concentrations of peptides exhibit increased G′. For KAFIL hydrogels, the G′ increases significantly in basic conditions.

**Figure S8.**
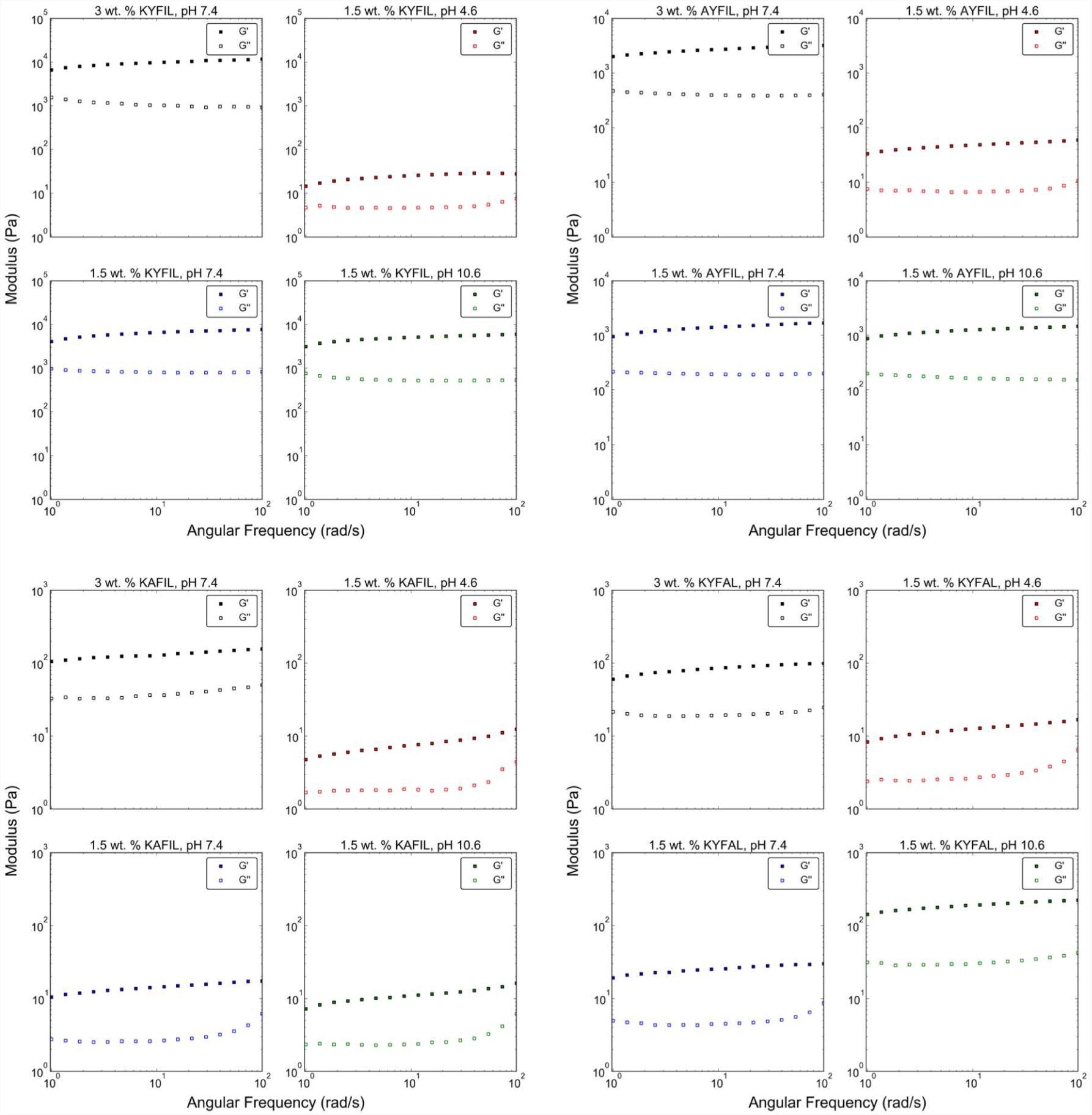
Frequency sweep of gelling pentapeptide sequences at constant strain at 0.1%. Measurements are carried out at 3 wt.% and 1.5 wt.%, and different pH conditions (4.6, 7.4, 10.6). For all sequences, G′ decreases significantly in acidic conditions. Higher concentrations of peptides exhibit increased G′. For KAFIL hydrogels, the G′ increases significantly in basic conditions.

**Figure S9.**
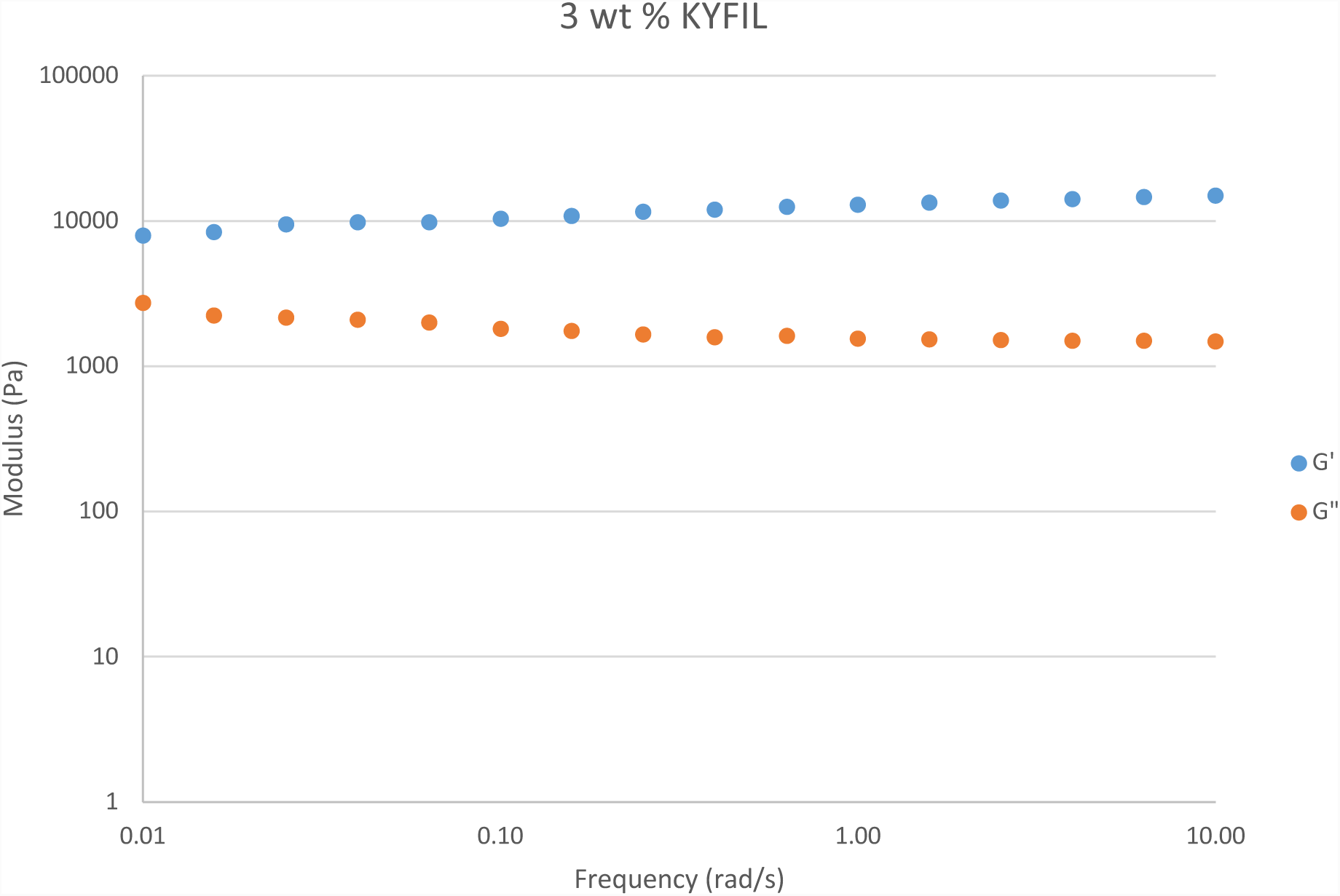
Frequency sweep of gelling pentapeptide sequence KYFIL at constant strain at 0.1%. Measurements are carried out at 3 wt.% from 0.01 to 10 rad/s to investigate the inherent dynamics of the hydrogel network.

**Figure S10.**
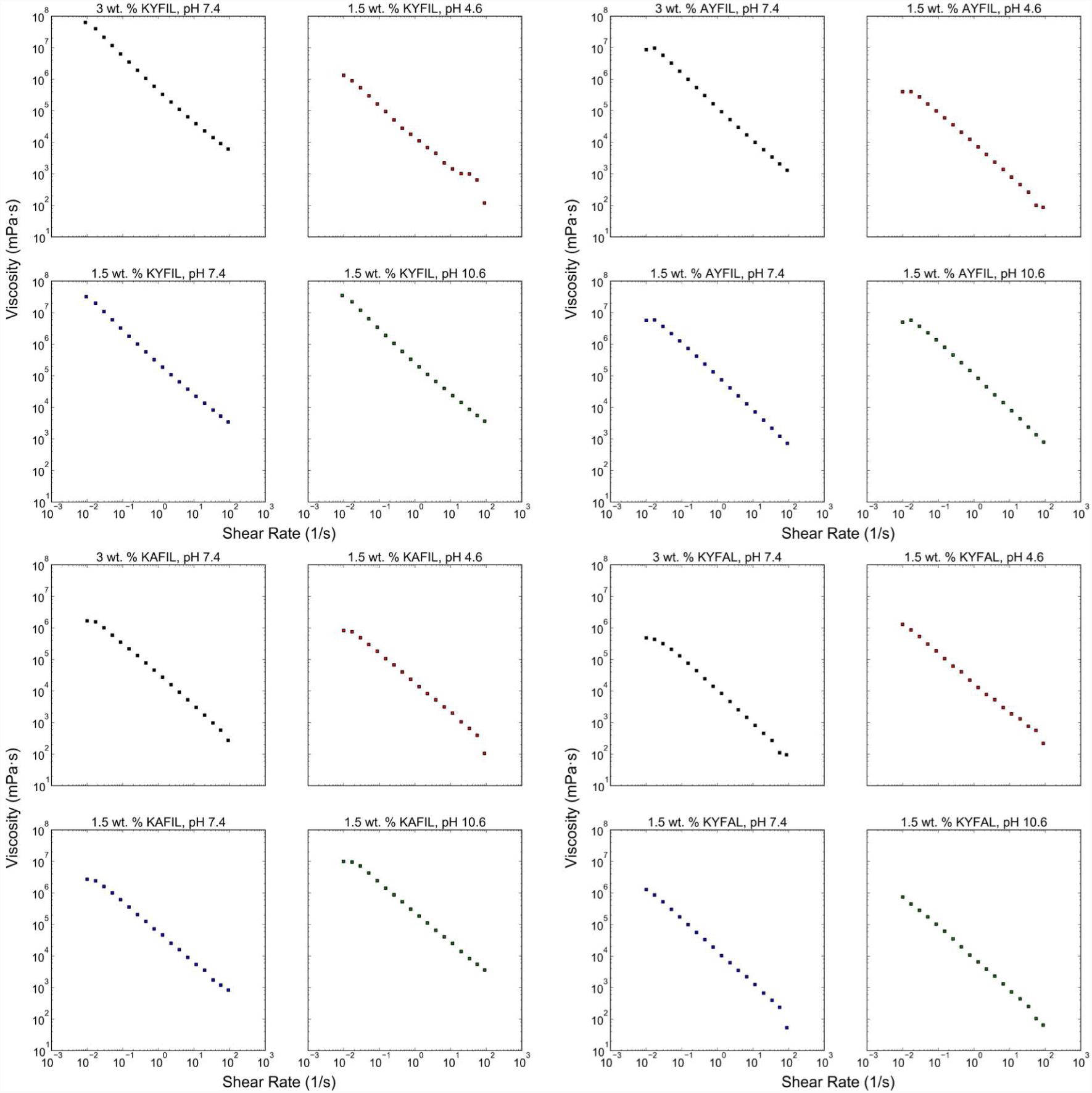
Apparent viscosity versus shear rate measurements of gelling peptide sequences at different wt.% and pH conditions. All hydrogels displayed shear-thinning behavior, in which the viscosity of each sample decreases with increasing shear rate.

**Figure S11.**
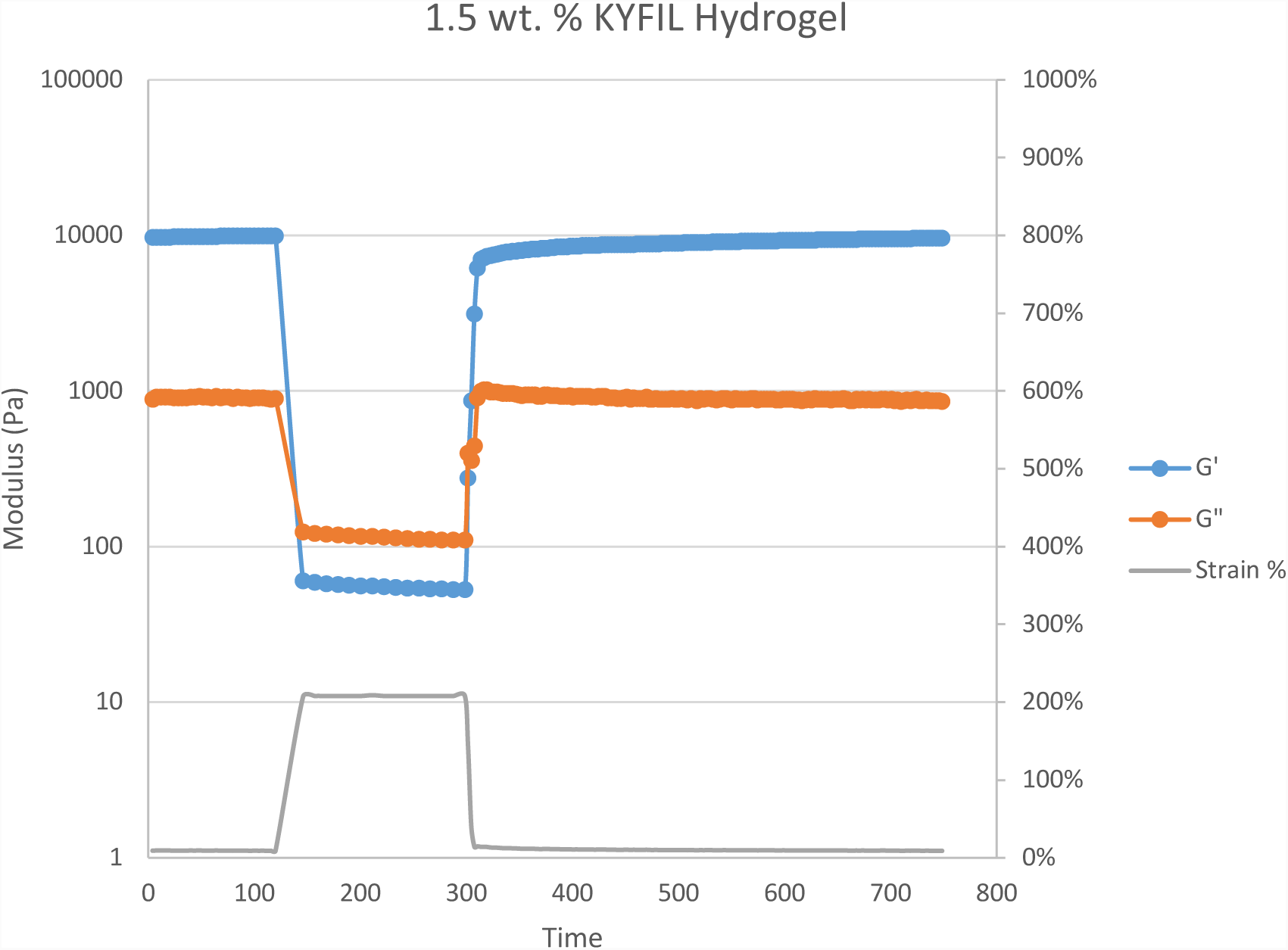
A thixotropy test was performed for 1.5 wt. % KYFIL hydrogels. A strain sweep of 0.1 % (100 s) followed by a 200 % strain (200 s), followed by a 400 s recovery period. The hydrogel is able to recover 90% of its initial G′ in 3.5 minutes, and 7 minutes to recover 96%.

**Figure S12.**
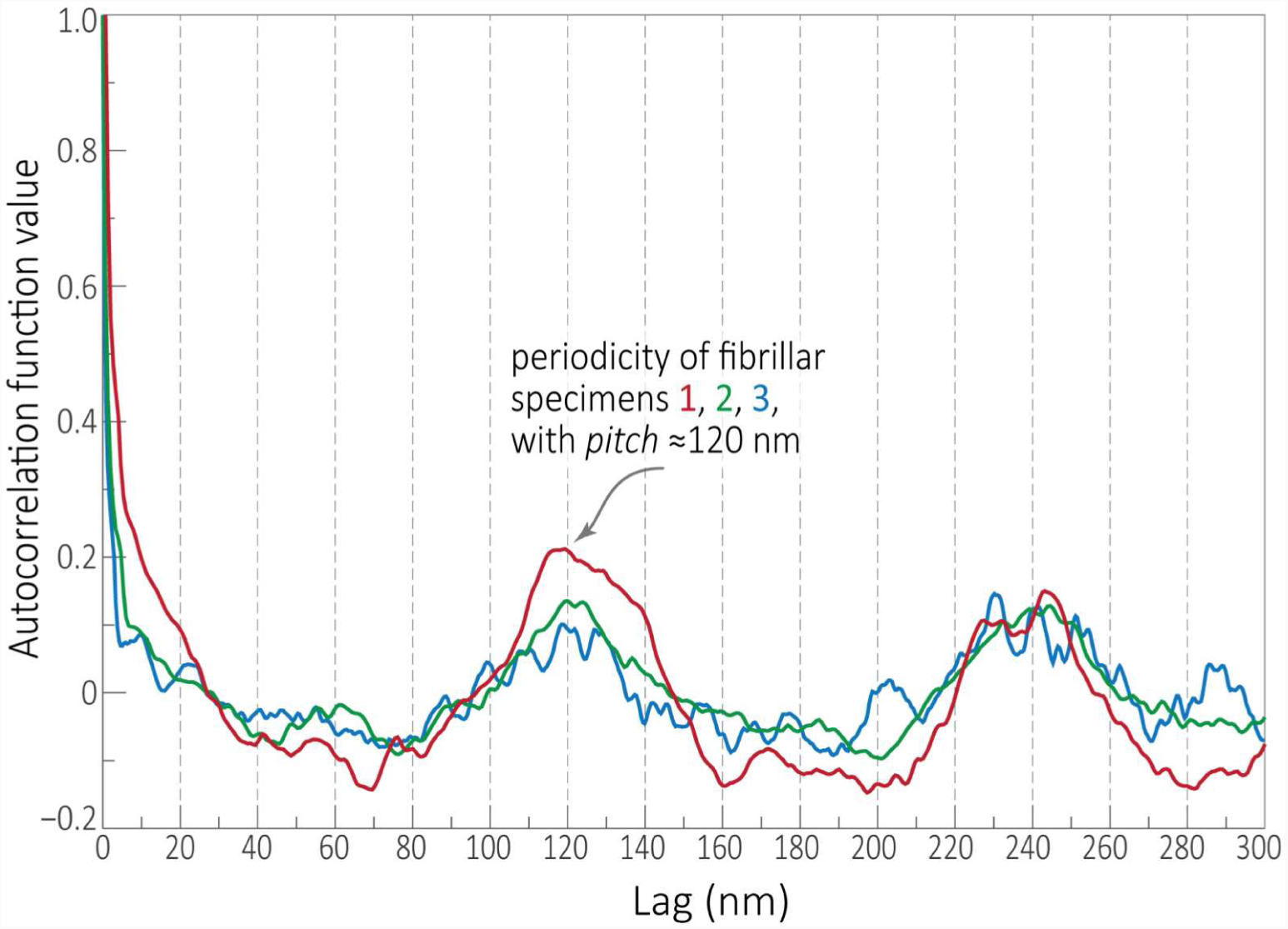
Periodicity of the fibrillar twist, as quantified by the intensity autocorrelation function (ACF). The ACF was computed for the height intensity of fibrils found in micrograph images of three independent TEM specimens (red, green and blue traces); the fundamental frequency can be seen to correspond to a distance (lag) of ≈120 nm.

**Figure S13.**
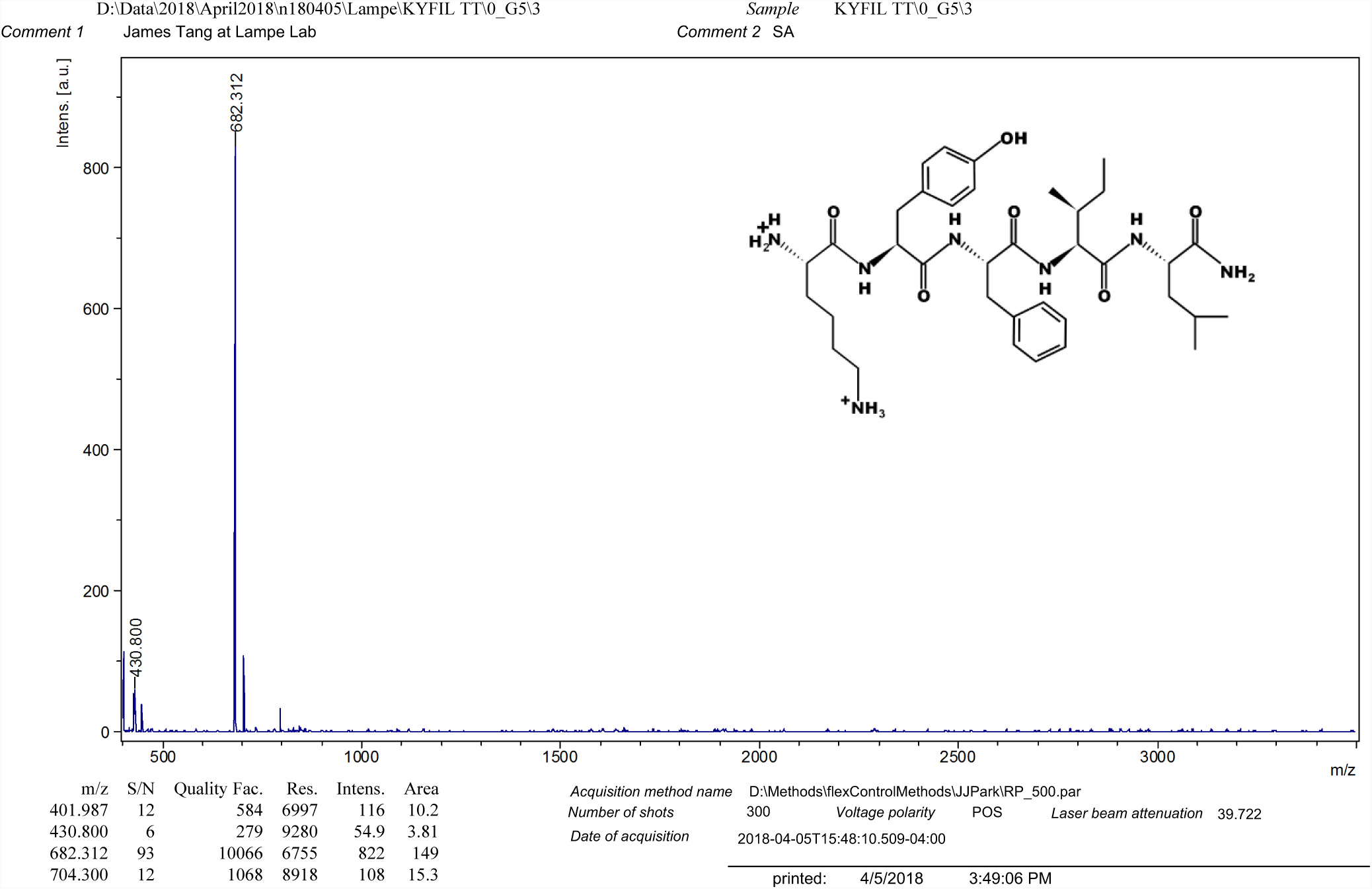
MALDI-TOF MS of KYFIL peptide. Expected mass [M+H^+^]^+^ = 682.87, observed mass = 682.312.

**Figure S14.**
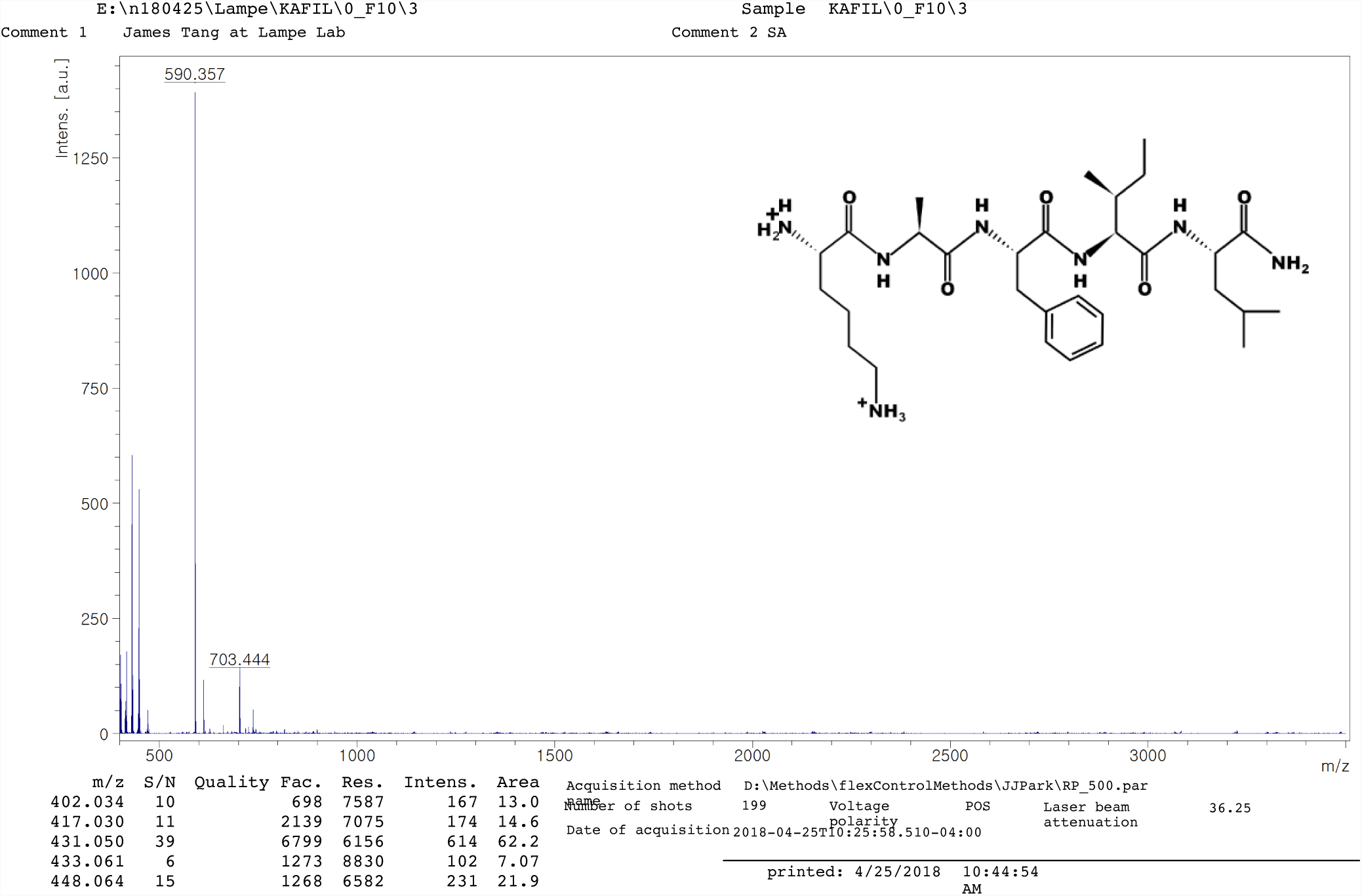
MALDI-TOF MS of KAFIL peptide. Expected mass [M+H^+^]^+^ = 589.77, observed mass = 590.357

**Figure S15.**
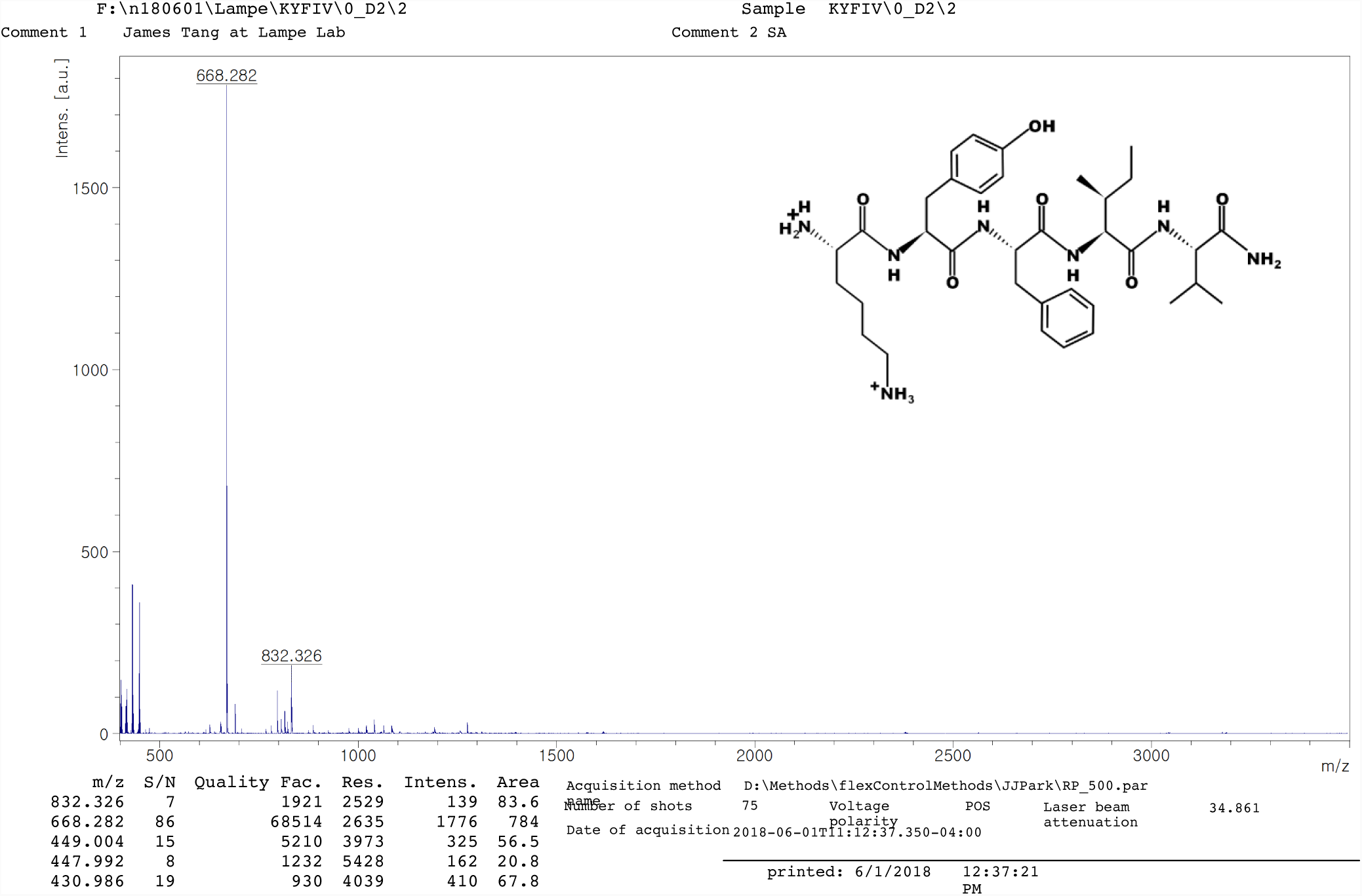
MALDI-TOF MS of KYFIV peptide. Expected mass [M+H^+^]^+^ = 667.84, observed mass = 668.282

**Figure S16.**
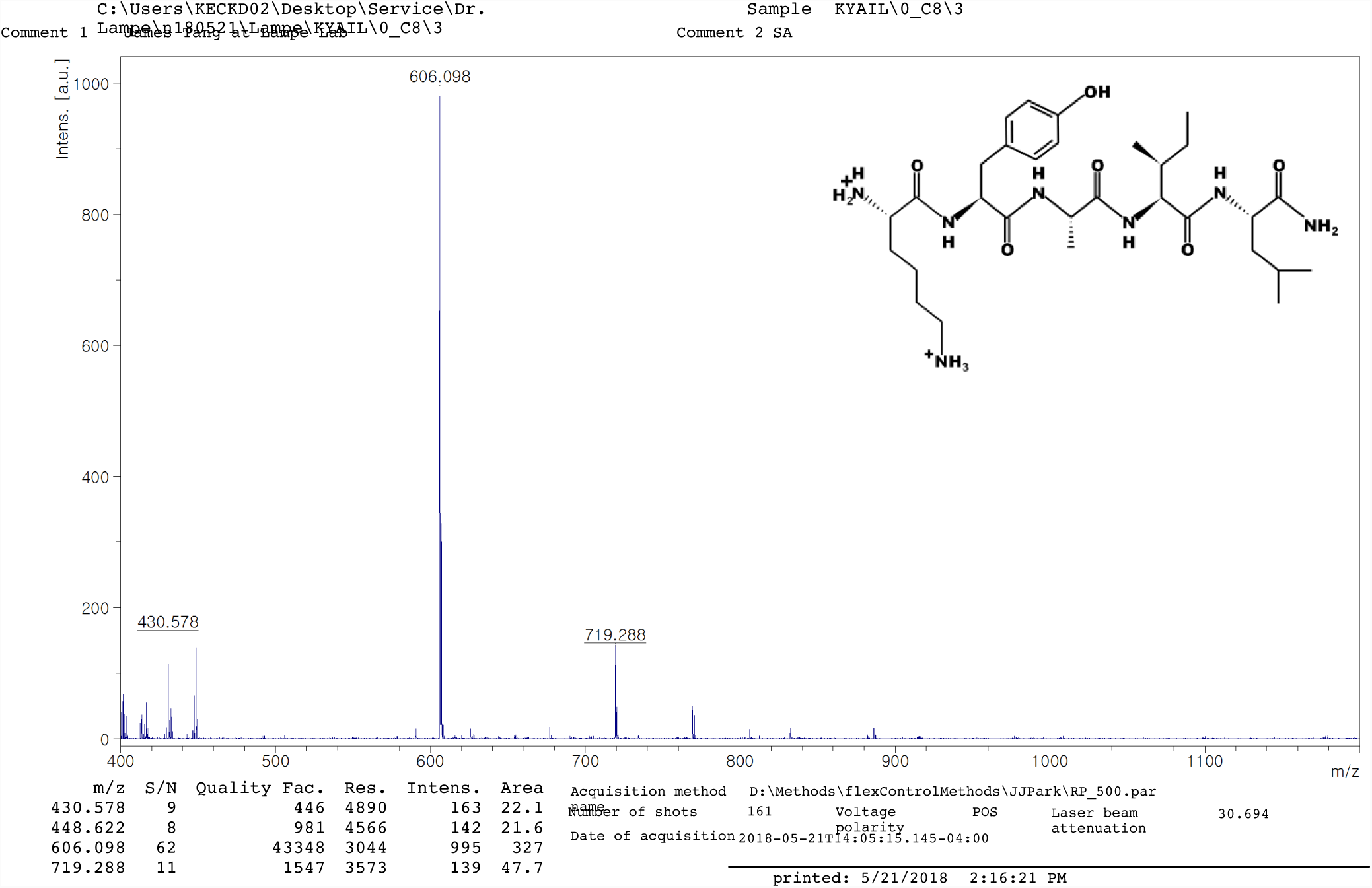
MALDI-TOF MS of KYAIL peptide. Expected mass [M+H^+^]^+^ = 605.77, observed mass = 606.098

**Figure S17.**
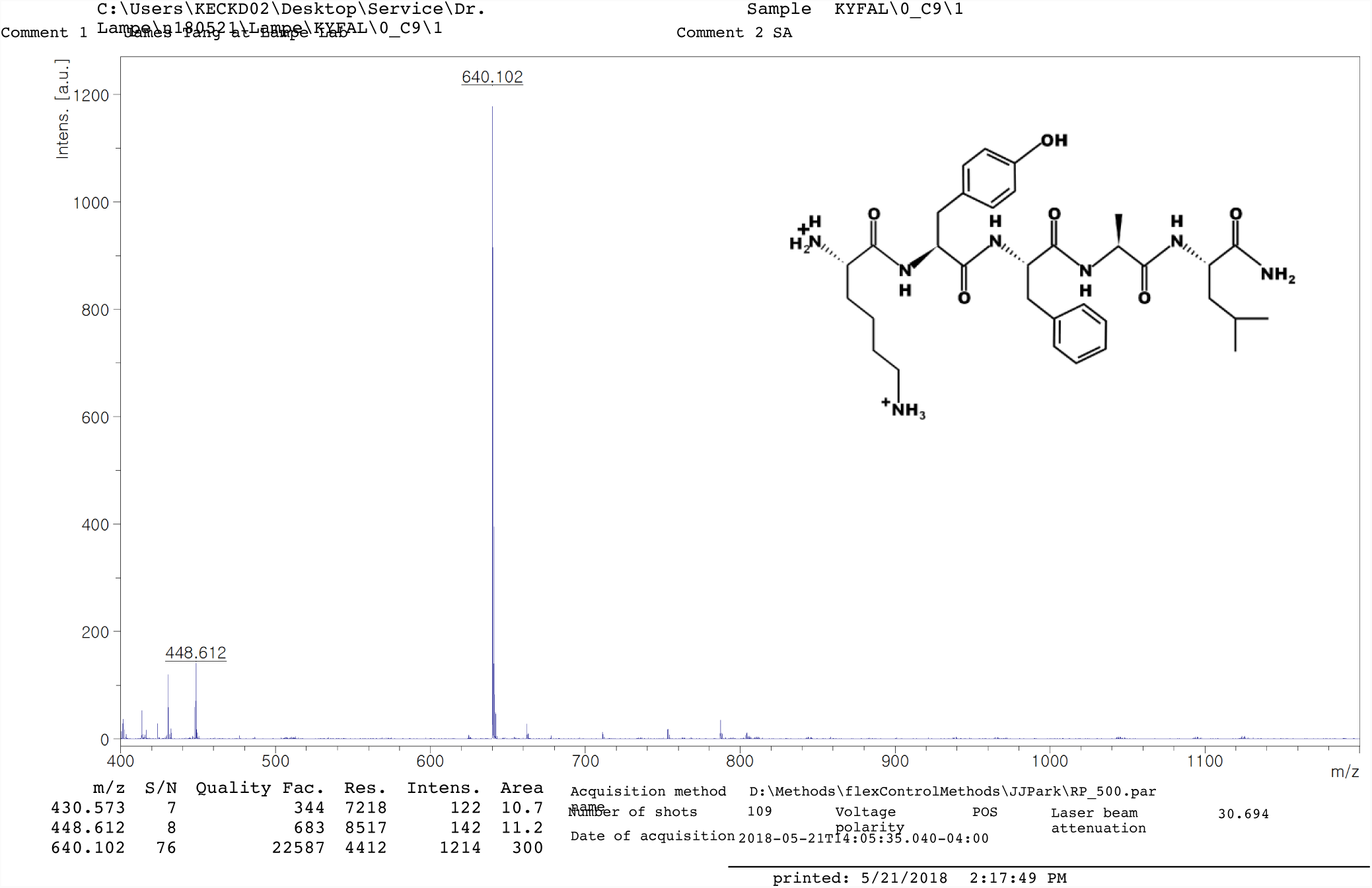
MALDI-TOF MS of KYFAL peptide. Expected mass [M+H^+^]^+^ = 639.79, observed mass = 640.102

**Figure S18.**
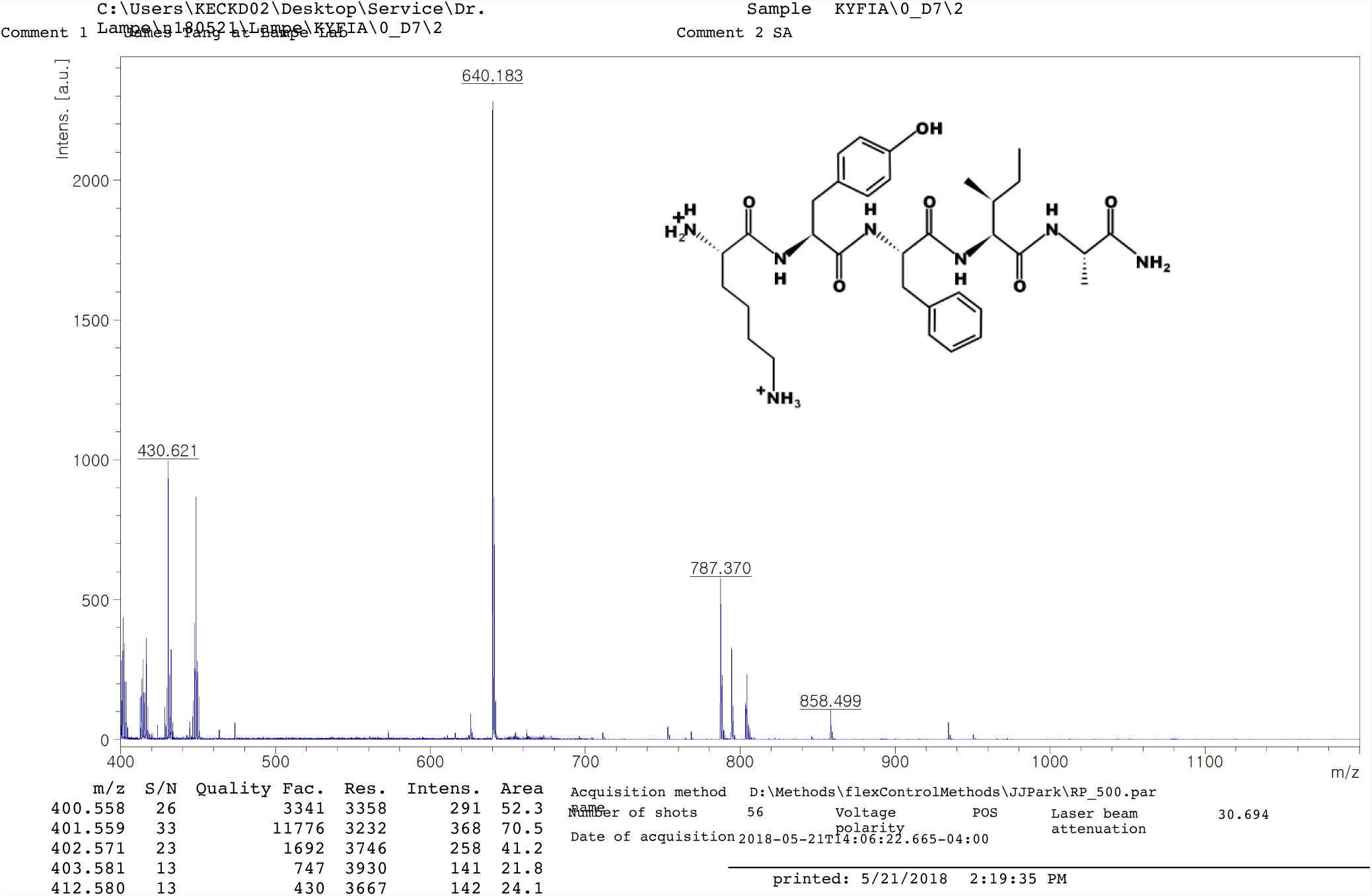
MALDI-TOF MS of KYFIA peptide. Expected mass [M+H^+^]^+^ = 639.79, observed mass = 640.183

**Figure S19.**
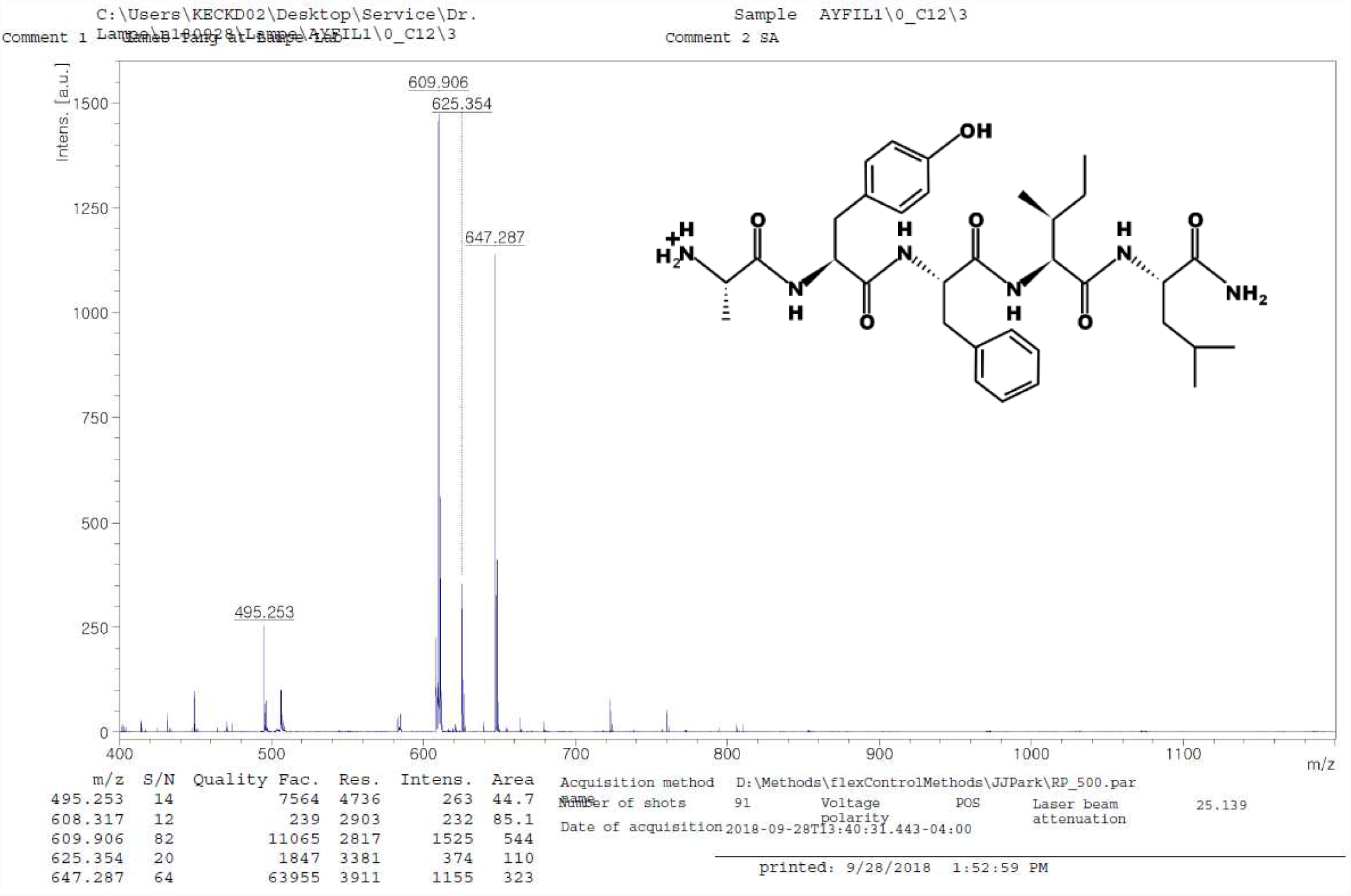
MALDI-TOF MS of AYFIL peptide. Expected mass [M+H+]+ = 624.77, observed mass = 625.354.

